# A model of neural population dynamics for flexible sensorimotor control

**DOI:** 10.1101/2025.05.15.654192

**Authors:** Hari Kalidindi, Frédéric Crevecoeur

## Abstract

Modern large-scale recordings have revealed that motor cortex activity during reaching follows low-dimensional dynamics, thought to reflect sensorimotor computations underlying muscle activation. However, the origin of these low dimensional patterns, and how they flexibly reorganize across different tasks remain unclear. Here we demonstrate that the key features of neural activity can naturally emerge in a linear model combining a random network with a biomechanical system. Remarkably, this model shows explicitly how a fixed network can achieve flexible control of multiple behaviours through optimal mapping of sensory feedback onto the network. Finally, analytical decomposition of the controller reveals that low-dimensional network dynamics follow directly from the propagation of low-dimensional feedback signals from the biomechanical plant through the network. This formalism provides a computational framework to interpret flexible motor control in the nervous system, which directly links neural population dynamics to sensorimotor behavior.

## Introduction

A central challenge in neuroscience is to explain how neural activity generates behavior and adapts to changing environments(Krakauer et al., 2017). To address this, recent advances have shifted focus from individual neurons to the collective dynamics of populations of neurons(Gao & Ganguli, 2015; Stringer & Pachitariu, 2024; Urai et al., 2022). Empirical studies show that the population dynamics can be effectively captured by rather low-dimensional, time-varying signals, referred to as *neural* factors (Churchland & Shenoy, 2024), or *latent dynamical state*(Gallego et al., 2017; Vyas et al., 2020). These factors have been interpreted as signals that the brain must generate to orchestrate behavior(Churchland et al., 2012; Churchland & Shenoy, 2024). Yet, fundamental questions remain: what is the relationship between neural factors and biological or physical variables that characterize the movement, and which factors are required for a given behavior?

Prevailing theories propose that neural factors emerge from recurrent connectivity among cortical and subcortical areas, forming attractors that shape population activity(Vyas et al., 2020). Artificial recurrent neural networks (RNN) capture this idea by developing stable patterns – such as tonic or multi-phasic activity – generating muscle commands when optimized on sensorimotor tasks(Aldarondo et al., 2024; Feulner et al., 2022; Guang et al., 2024; Gurnani et al., 2024; Kalidindi et al., 2021; Lillicrap & Scott, 2013; Michaels et al., 2020; Saxena et al., 2022; Sussillo et al., 2015). However, generalization to other tasks remains challenging for these models, which rely on training based on novel relevant datasets(Zador et al., 2023). Their behavior is often difficult to interpret due to non-linearities and dependency on both training data(Zador et al., 2023), and training algorithms(Codol et al., 2024). This raises the question: are simpler, more interpretable models sufficient to account for some key features of motor cortical dynamics?

In contrast to trained artificial networks, primates exhibit remarkable flexibility as they can rapidly adapt to novel contexts, or even update control within an ongoing movement(Braun et al., 2009; Crevecoeur, Mathew, et al., 2020; Kalidindi & Crevecoeur, 2023; Lake et al., 2016; Saxe et al., 2021). Although development certainly forms a wide range of behaviors akin to long term training, recent computational modeling studies suggest that contextual inputs to the network can evoke patterns suitable to a given task(Beiran et al., 2023; Kao et al., 2021; Lindén et al., 2022; Logiaco et al., 2021; Mante et al., 2013; Remington et al., 2018; Schimel et al., 2023; Sohn et al., 2021; Stroud et al., 2018). However, it remains unclear how inputs are computed and adjusted across diverse sensorimotor tasks, such as different instructions or loading conditions.

Here we propose a unifying theoretical framework that explains both the emergence of low-dimensional factors in motor cortical dynamics, and how inputs to the network should be computed to accomplish the task at hand. We developed a linear model coupling a random network to a biomechanical limb and simulated various tasks using stochastic optimal control(Åström, 1970; Schimel et al., 2023; Todorov & Jordan, 2002). In this framework, the feedback control policy maps the feedback about the state of the whole system (network and limb) onto optimal network activations. We demonstrate that this approach directly links the parameters of the controller to the network activity across tasks without the need for example-driven (re)training. Remarkably, our model reproduced key features observed in motor cortex, including preparatory activity anticipating the upcoming movement(Churchland et al., 2010; Kaufman et al., 2014), rotational dynamics during movement(Churchland et al., 2012; Sabatini & Kaufman, 2024), and activity during movement preparation and execution periods restricted to orthogonal subspaces(Elsayed et al., 2016). Furthermore, we show that this linear formulation offers an novel view on the origin of low-dimensional factors using the low-rank hypothesis of complex systems(Thibeault et al., 2024): although the network connectivity is high-dimensional, its activity is mainly driven by inputs carrying sensory feedback and task parameters, yielding low-dimensional patterns of activation. Together, our results propose a formal link between sensorimotor control and low-dimensional population activity for a fixed random network, offering a powerful candidate framework to interpret the neural basis of flexible motor control.

## Results

We constructed a super-system composed of a randomly connected recurrent network (*n* = 100 nodes) coupled to a limb (Fig. 1a). The network activity, ***r***(*t*) ∈ ℝ^100^, was represented as a standard rate equation(Kalidindi et al., 2021; Kao et al., 2021; Schimel et al., 2023; Sussillo et al., 2015). The network output controlled one of the two limb models: a point-mass moving in a cartesian plane or a two-link planar arm. Both models featured two-dimensional actuation, with cartesian forces applied along the plane for the point-mass, and joint torques applied to the shoulder and elbow joints for the planar arm. The limb state, ***q***(*t*) ∈ ℝ^66^, encompassed position, velocity and actuation forces/torques. Full details of the network and limb dynamics are provided in the Methods. For compact representation, the linear dynamics governing the network-body system were jointly described by:

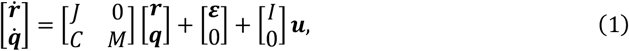

where *J* ∈ ℝ^100×100^ was the random state transition matrix, ***ε*** ∈ ℝ^100^ was a constant input vector inducing heterogenous spontaneous activity, *C* was the random linear readout that mapped the network activity to limb actuation, and *M* was the state transition matrix of limb dynamics (the explicit dependency on time was omitted from Eqn. 1 for clarity). The control input ***u***(*t*) ∈ ℝ^100^ influenced only the network activity. The identity matrix *I* denotes that each control input component activated a single network unit.

**Figure 1.**
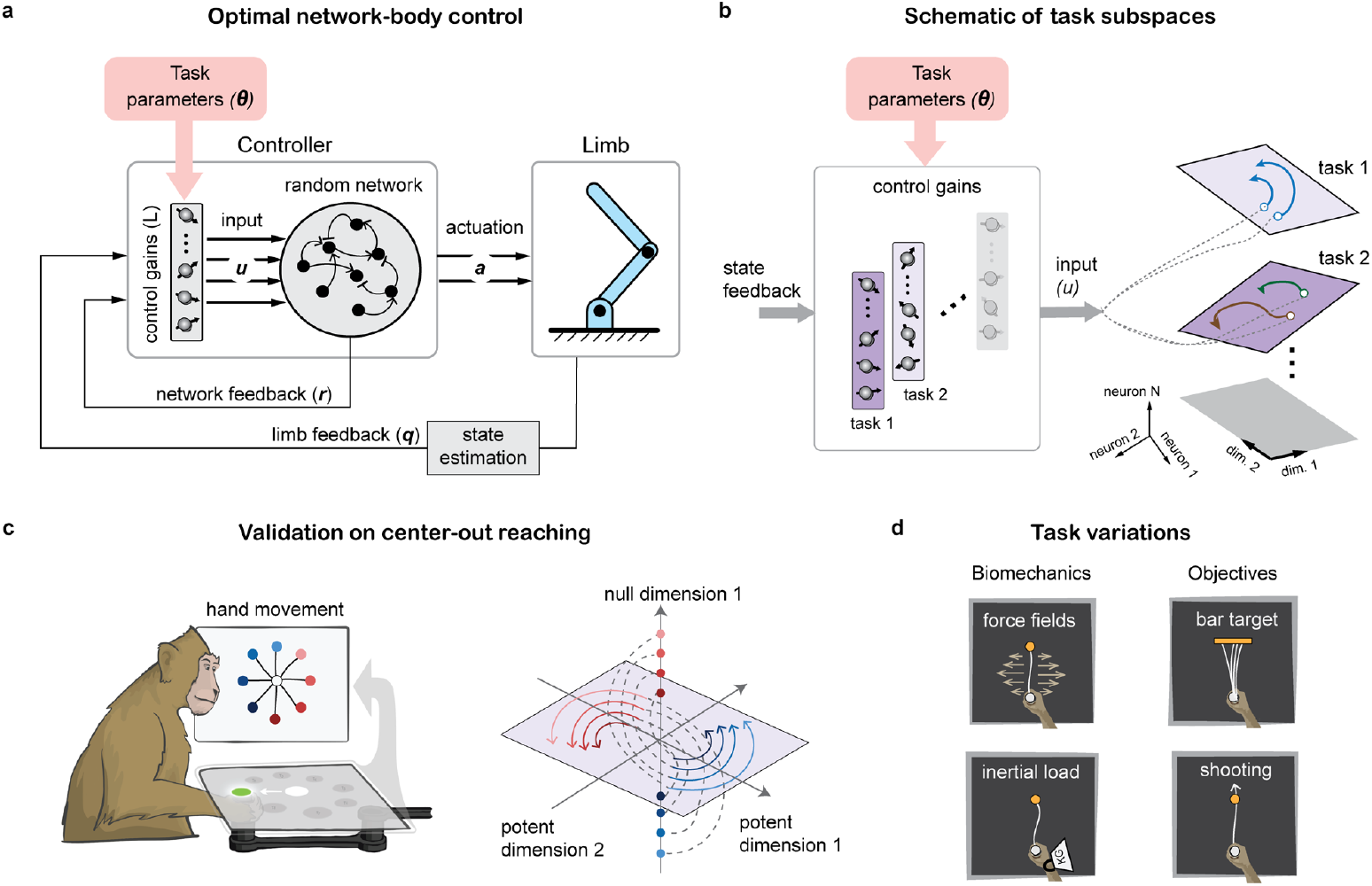
Schematic representation of the network-body system. a) Task-dependent control of network-body system through tunable control gains. b) illustration of how the tunable control gains may reshape the low dimensional neural trajectories based on task parameters. c) Illustration of a stereotypical prepare-to-reach task, showing how the neural population activity is driven into orthogonal subspaces for preparation and movement, where dots represent preparatory states, curves represent activity over time and color coded to different directions of reaching d) Selected benchmark tasks taken from standard motor control literature used to test the malleability of sensorimotor control in primates dependent on changes in task objectives (flexibility) and biomechanics (adaptation).

The objective was to find inputs ***u***(*t*) that steer the network activity to make the limb move optimally as defined by a cost-function. We adopted the formalism of receding-horizon to allow modelling changes in task instructions such as the unexpected occurrence of a “GO” cue presented during movement preparation (see Methods). This problem was solved as follows: we determined the control inputs that minimized the expected cost (𝒥(***q, u***, *t*)) over a time-horizon Δ*h* defined by:

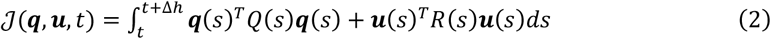

We then applied the first time-step of the control law and repeated the process by moving the time-horizon forward. Here, *Q*(.) is a positive-semidefinite matrix penalizing deviations from desired limb states (note that state vector includes the target, omitted here for simplicity, see Methods), and *R*(.) is a positive-definite matrix penalizing large network inputs (***u***).

We now simply consider the structure of the controller to interpret its effect on the network-body dynamics across task variations. For the given system dynamics (Eqn. 1) and quadratic cost function (Eqn. 2), the optimal control inputs for each time-horizon can be computed using the classical Linear Quadratic Regulator (LQR) approach(Åström, 1970; Todorov, 2005) (see Methods), which yields an optimal state-feedback control law that can be decomposed as follows:

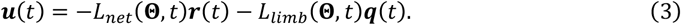

The set **Θ** encompasses all the system and cost parameters (i.e. **Θ** = {*Q, R, J, C, M*}), and *L*_*net*_ and *L*_*limb*_ are the control gains applied to the network and limb states, respectively. Substituting this feedback control law (Eqn. 3) into the system dynamics (Eqn. 1) gives the closed-loop system:

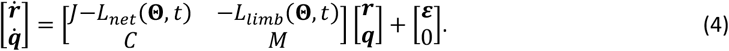

Rewriting the system in this closed-loop form offers an interpretation of the controller structure: this optimal control scheme enables multiple tasks (represented by **Θ**) by dynamically adjusting the coupling of network nodes (*J* became *J*−*L*_*net*_(**Θ**, *t*)), and the feedback about the limb states through the matrix −*L*_*limb*_(**Θ**, *t*) (Fig. 1a). In essence, this approach leverages the linear structure of the system to compute the optimal network connectivity suitable for a given set of parameters without data-driven (re)training. Thus, we may expect that different tasks (and their corresponding sets of parameters) induce different network activations, resulting in various shapes and trajectories of the network states representing neural population activity (Fig. 1b). We show below that this model remarkably reproduces previously published properties (Fig. 1c), and makes quantitative predictions of neural activities associated with standard tasks (Fig. 1d).

### Emergence of low-dimensional factors for preparation and execution

We first validate the model by showing that it reproduces hallmark features of monkey’s primary motor cortex (M1) activity during a standard center-out reaching task(Churchland et al., 2012; Elsayed et al., 2016) (Fig. 1c). Specifically, using a receding-horizon control scheme(Guigon, 2023; Maciejowski, 2002) (see Methods), the model generated realistic hand trajectories in a delayed prepare-to-reach paradigm, including bell-shape velocity profiles following the GO cue (Fig 2a). At the level of individual network nodes, we observed heterogenous activity patterns, with tonic levels of activity during the preparatory phase that quickly encoded the impending target, and a subsequent phasic activity linked to movement execution (Fig 2b). Both the amplitude and the phase of individual responses varied across nodes and movement directions. Notably, many nodes exhibited an overshoot immediately after target onset, resembling M1 recordings(Kaufman et al., 2014; Lara et al., 2018). We observed that this overshoot occurred due to the cost parameters, as increasing the penalty on control inputs reduced it (simulations not shown).

**Figure 2.**
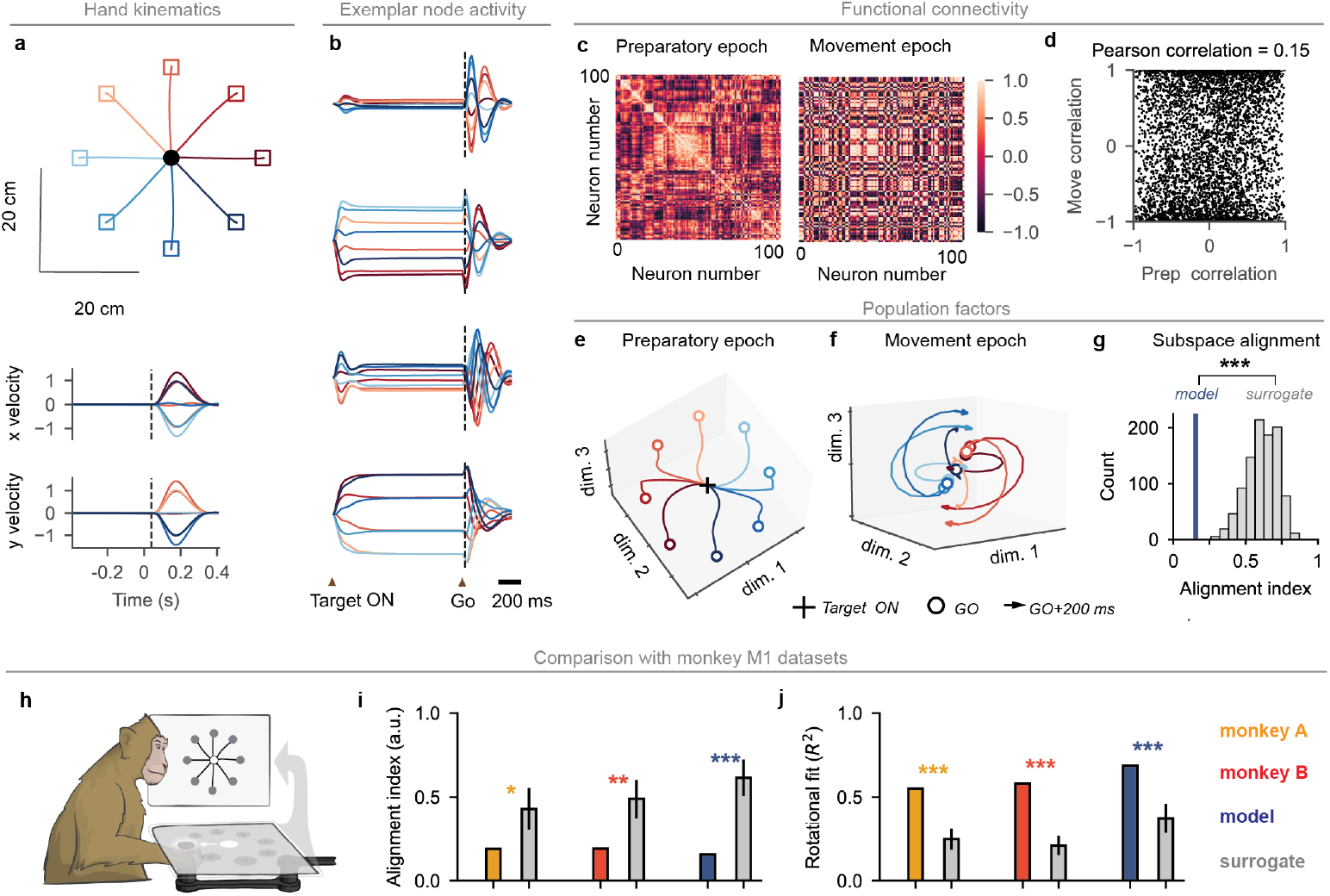
Feedback control produces meaningful behavior and reproduces M1-like preparation and movement-related “factors”. a) Hand kinematics for the simulation of a point mass system b) exemplar neural firing rates from selected units, observe that the activity of some neurons depends on the target of the pending movement. c) Pairwise correlations in preparatory and movement periods d) Linear relationship between pairwise-correlations from movement to preparatory period. The lack of pattern shows that activities from one epoch is statistically independent of that of the other epoch e) neural activity along the principal modes in the preparatory period. f) same as (e) for the movement period. g) alignment between preparatory and movement period activities. The histogram shows distribution of alignment indices across surrogate data and the vertical line represent the alignment of subspaces from the principal modes of preparatory and execution epochs (low alignment corresponds to orthogonal) h) comparison of the model with monkey datasets, monkey A(Churchland et al., 2012), monkey B(Suresh et al., 2020) i) comparison of alignment index of top six preparatory and movement period PCs j) The goodness of fit (R^2^) obtained by fitting the top six PC trajectories to a rotational dynamical system using the jPCA procedure. Asterisk symbols indicate the p-values from one-tailed test: ^*^p-val < 0.05, ^**^p-val < 0.01, ^***^p-val < 0.005

To characterize the interactions between nodes across epochs, we first analyzed functional connectivity by computing pairwise correlations of time-series activity during the preparatory (150 to 50 *ms* before the GO cue) and movement epochs (50 to 150 *ms* after the GO cue). Fig. 2c shows that the average pairwise-correlations across targets observed during preparation was not maintained during movement. Consequently, the pairwise-correlations among nodes during preparation were not linearly related to those observed during movement. This resulted in a low Pearson correlation-coefficient between epochs (Fig 2d) indicating a reorganization of network interactions driven by task objective to prepare or execute movement.

To examine the dominant features of population activity and link them to previously reported neural factors, we performed principal component analysis (PCA) on the network activity during preparatory and movement epochs (see Methods). The dominant principal components (PCs) in the preparatory epoch encoded target information prior to movement onset (Fig. 2e). This finding demonstrates that directional encoding of the upcoming movement occurred within a subspace of ℝ^*n*^ that did not produce any motor output. This observation is consistent with experimental observations where neuronal activity unfolds along “output-null” dimensions that do not produce any movement(Churchland & Shenoy, 2024; Elsayed et al., 2016). The same PCA applied to the population activity after the GO cue revealed rotational trajectories (Fig. 2f), closely resembling rotations reported in M1(Churchland et al., 2012). Notably, the control scheme showed distinct responses between preparatory versus movement epochs because the task objectives (*Q*) changed: during preparation deviation from start was penalized, whereas during movement deviation from the goal was penalized. As a result, the controller allowed the same state-feedback signals to produce different network trajectories depending on whether the system was preparing or executing a movement.

This task-dependent control scheme also captured another well-known feature of M1 activity: the orthogonality between preparatory and execution related activity(Elsayed et al., 2016). Applying PCA to network activity in each epoch separately, the preparatory-PCs explained little of the variance observed during the movement, and vice-versa. To quantify the degree of overlap, we computed the alignment index(Elsayed et al., 2016) (see Methods), measuring how much variance in preparatory epoch was explained by the first six *PCs* from the execution epoch, normalized by the variance explained by the first six *PCs* from the preparatory epoch. When the activities during preparation and execution phases are confined to orthogonal output-null and output-potent subspaces respectively without any overlap, the alignment index is zero. In our model, the network activities exhibited a low alignment index, consistent with recordings from M1 (Elsayed et al., 2016), and significantly lower than surrogate datasets generated via the tensor maximization entropy (TME) procedure (Fig. 2g, see Methods)

The foregoing observation highlighted low alignment between preparatory and movement related activities, as well as rotational trajectories. To evaluate the model’s resemblance to the motor cortex quantitatively, we compared its behavior with publicly available datasets of neural recordings in monkeys (Fig. 2h)(Churchland et al., 2012; Suresh et al., 2020). Specifically, we compared the alignment index and rotational dynamics in both the model and M1 data. Both the model and M1 recordings exhibited low alignment indices, substantially lower than those observed in surrogate datasets (Fig. 2i). Rotational dynamics were quantified using jPCA method(Churchland et al., 2012), which was used to fit the top six principal components to a constrained dynamical system with purely rotational structure (see Methods). The goodness of fit, measured by *R*^2^, was consistent across M1 recordings and model simulations, and significantly larger than surrogate datasets generated via the TME procedure (Fig. 2j).

To further characterize the rotational structure from the firing rate without dimensionality reduction, we computed the complex-valued gyration number(Kuzmina et al., 2024). The real and imaginary parts of the gyration number capture the normalized decay and rotational components in the population activity, respectively (see Methods). The model and M1 recordings exhibited comparable levels of rotation and decay, whereas shuffled datasets showed markedly reduced rotational component (Supplementary Fig. 1).

Thus, closed-loop control of the network-body system naturally gave rise to low-dimensional neural trajectories in orthogonal preparatory and movement subspaces, with rotational population dynamics closely paralleling patterns of neural activities recorded in M1. Remarkably, these features emerged from a linear system combined with a quadratic cost. No additional constraint was required, such as enforcing simple network dynamics(Sussillo et al., 2015), or minimizing trajectory tangling(Russo et al., 2018; Saxena et al., 2022) as expressed in previous models.

We made several additional observations that are reported in the Supplementary Material, which can be summarized as follows: our results appear robust against variations in the model structures for comparable networks and inputs. Supplementary Fig. 2 shows that the results for the center-out reaching task were similar when coupling the network to a two-link arm instead of a point-mass as shown in Fig. 2. Then, we also observed low alignment between preparatory and movement related activity, as well as rotations, in fully connected recurrent networks without any excitation-inhibition balanced structure, and randomly sampled Erdös-Rényi networks (see Methods and Supplementary Fig. 3, 4).

To gain further insight into the model’s properties, we examined the effect of state feedback control on the network dynamics. First, spectral analysis of the state transition matrix revealed that the distribution of eigenvalues contained complex pairs of eigen values associated with oscillatory behavior (Fig. 3a). In the initial, randomly configured networks, many eigenvalues exhibited large absolute imaginary components corresponding to high-frequency oscillations, and real parts close to imaginary axis, indicating slow decay. Interestingly, the closed form expression (*J*−*L*_*net*_) modified the distribution by reducing the maximum absolute imaginary part of the eigen values and decreasing the spectral abscissa (Fig. 3a, b).

**Figure 3.**
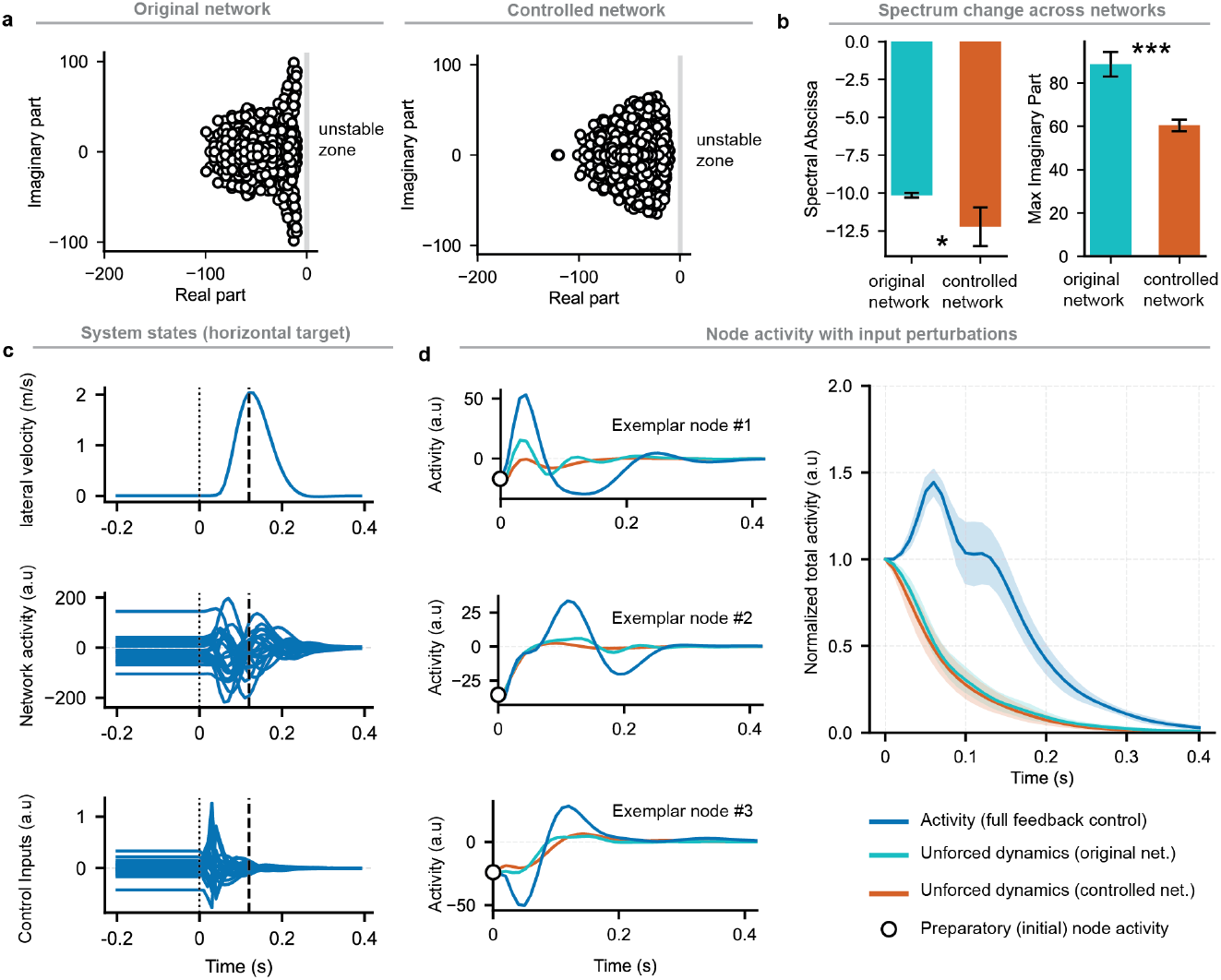
Model analysis. a) Eigen value spectrum of an exemplar original network (left) and the same network when modified by the state feedback control law (right, Eqn. 4). b) Change in spectral abscissa (maximum real part) and in the maximum imaginary part of the eigen values across 5 networks c) Simulated hand velocity, network activity and control inputs while reaching to the target aligned with the x-axis. Dotted and dashed vertical lines represent the time of GO cue and time of maximum velocity respectively d) Left panels: Node activity with full state feedback inputs compared with the unforced dynamics after the same initialization. Right panel: Normalized total activity across nodes and reach directions. ^*^p-val < 0.05 ^***^p-val < 0.005 from paired two-sample ttest across N=5 samples.

Next, we analyzed the role of state feedback inputs in driving the network activity. After the GO cue, inputs displayed a smooth, time-varying profile that persisted beyond peak velocity (Fig. 3c). Interestingly, input signals (***u***(*t*)) decayed faster than the network activity, suggesting that movement execution was supported by a sustained activation generated by the network’s intrinsic dynamics. To evaluate its contribution, we examined both the unforced dynamics of the initial, randomly configured network (*J*) and the dynamics of the controlled network (*J*−*L*_*net*_) after setting the input to zero following the ‘GO’ cue. The activity of individual nodes revealed that the unforced dynamics were characterized by high-frequency, low-amplitude oscillations that decayed more quickly than the feedback-driven activity. We quantified the normalized total activity across nodes and reach directions, scaled to the initial preparatory state. Networks with state feedback inputs showed a rise in activity followed by a slow decay, whereas unforced dynamics exhibited a much faster decay (Fig. 3d). These results demonstrate how state feedback inputs interact with intrinsic network dynamics to shape the network activity during reaching movements.

### Variation in movement dynamics

Building on the validation of the model in the center out reaching task (as shown in Fig. 2), we now illustrate its predictive power. Indeed, we can change any task parameter value (i.e. change a subset of **Θ**) and interrogate the model about the resulting neural activities. Using paradigmatic examples from motor control literature (Fig. 1d), we demonstrate how task parameters shape neural trajectories, enabling quantitative predictions about neural population dynamics across a range of tasks. Two subsets of task parameters are often studied: (1) the task objectives (the weighting matrices *Q* and *R* penalizing states or inputs in the cost function, expressed in Eqn. 2), and (2) the system dynamics (the matrix *M*). Based on the fact that the control gains, and hence the network inputs, are modulated as a function of the above-mentioned task parameters (Eqn. 3), we can explore how these parameters shaped the network dynamics.

We first examine the influence of velocity-dependent force-fields, widely used to study motor adaptation(Thoroughman & Shadmehr, 2000), on the network dynamics. A velocity-dependent force-field of the form 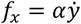 and 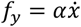 modifies the limb dynamics by coupling forward speed with lateral force and lateral speed with forward force. Assuming that the controller has knowledge of this force field, the model was able to produce straight hand paths towards the target after the GO signal (Fig. 4a), countering the applied lateral forces with scaled hand forces (Fig. 4a, bottom-right). Note that we do not model the process of adaptation, but we assumed that the controller knows the force-field parameter. Although no movement occurred prior to the GO cue, a few individual nodes encoded the level of the force-field parameter (*α*) during the preparatory phase, without generating motor output (Fig. 4b). Hence, network activity in the output-null space already encoded the strength of force field parameter. This was followed by a scaling in the amplitude of rate responses in individual nodes during the movement epoch (Fig. 4b).

**Figure 4.**
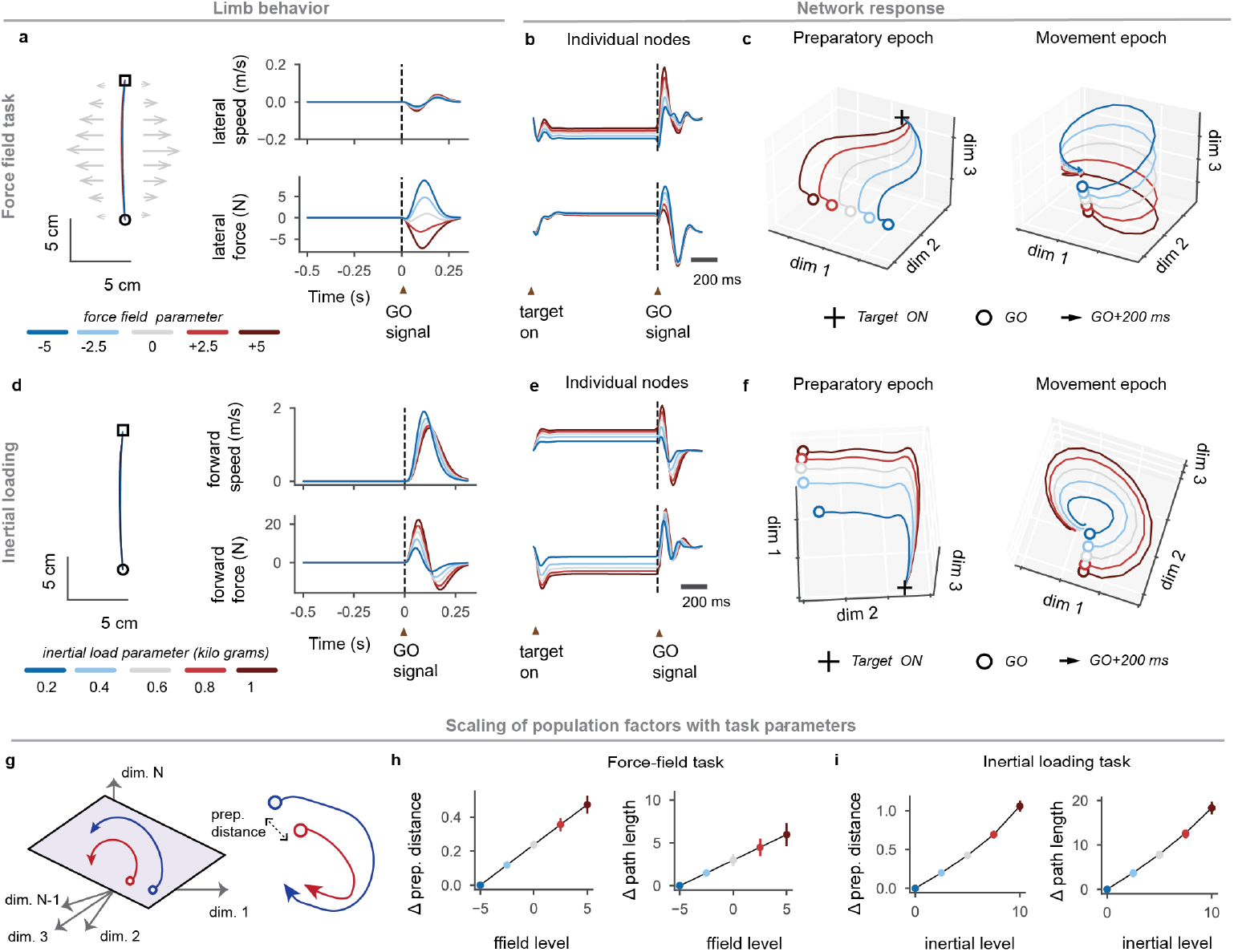
Flexible control to changes in biomechanical requirements. a) representation of the task with clockwise or counterclockwise force field of different strength, and hand kinematics with varying levels of lateral force applied to counter the force field. b) Exemplar neuron firing rates c) projection of the network trajectories in the principal mode of the preparation and movement epochs for five different levels of orthogonal force-field parameter. (d-e-f) same as above for five different levels of limb inertia. g) Illustration of the key measures of variation in neural factors: h) change in preparatory and movement related factors for force-field task. i) Same as h for inertial loading task. The circle markers depict mean across N=5 networks, and the error bars indicate the s.e.m.

At the population level, the dominant PCs encoded force field strength during the preparatory epoch, followed by rotational trajectories in the movement epoch (Fig. 4c). Notably, these rotations occupied different planes but maintained a consistent direction even when the force field was applied in opposite directions (−5 to +5).

Next, we quantified the changes in neural population responses across force-fields by computing the path changes compared to the baseline parameter level (*α* = 0) (Fig. 4g). The change in preparatory state was measured as the distance between the preparatory state at the GO signal and at the baseline force-field parameter level. Movement-related changes in PCs were quantified as the sum of squared differences between points on the PC trajectory and the baseline trajectory. Across different networks, both preparatory and movement period PCs scale with the strength of the force-field parameter (*α*) (Fig 4h). Indeed, analyzing the corresponding control gains across varying force-field values revealed that these gains changed with *α* during both preparatory and execution epochs, thereby shifting network activity into different preparatory states that scaled with the force-field strength even when moving to the same spatial target.

We next asked whether this modulation in neural trajectories extended to other changes in limb dynamics. In a second set of simulations, we varied inertial loads by simulating different masses of the hand while reaching to the target (Fig. 4d). Similar to the force-field task, individual nodes encoded the inertial load level *m*_*load*_ (Fig. 4e) during both preparatory and movement epochs. Applying PCA again showed that the dominant PCs in each epoch reflected the load magnitude (Fig. 4f). As before, both preparatory and movement PCs scaled with the level of loading (Fig. 4i), confirming that parameters altering limb dynamics produced feedback modulation that reshaped the network activity.

To assess the model properties with multiple targets and loading conditions, we simulated reaching tasks involving eight spatial targets under varying velocity-dependent force fields and inertial loads (Supplementary Fig. 5a). In both conditions, preparatory principal components (PCs) exhibited affine encoding: preparatory states varied with the magnitude of the load (*m*_*load*_) or force field parameter (*α*), while offsets were determined by the target location (Supplementary Fig. 5b). Notably, the radial organization shown in preparatory states is consistent with the empirical evidence from motor adaptation studies(Sun et al., 2022; Vyas et al., 2018). Under force field conditions, preparatory shifts occurred tangentially (toward or away from adjacent targets), whereas inertial loading induced radial scaling of preparatory states. During movement, PCs traced rotational trajectories that varied systematically with both target direction and mechanical loading (Supplementary Fig. 5c), indicating an interaction between spatial goals and limb dynamics. Quantitatively, increasing the angular separation between target directions was related to greater distance between preparatory states in the low dimensional, projected space (Supplementary Fig. 5d). Overall, varying the limb dynamics led to a combination of scaling and directional variations in the PCs of network population activity.

### Variation in task objectives

We now use the same approach to show a modulation of population activity in response to variation in task objectives (varying *Q* matrix in Eqn. 3). To test this, we simulated two tasks typically used in human behavioral experiments: the bar task(Gallivan et al., 2016; Nashed et al., 2012), and a shooting (or a hit) task(Česonis & Franklin, 2022).

In the bar task, the hand must reach a target with different levels of penalty on the lateral endpoint deviation, leading to variation in the lateral end position (Fig 5a). When a brief mechanical load was applied to perturb the movement laterally, the model corrected errors more vigorously in directions where errors were more costly (high lateral cost), leading to smaller deviations and higher corrective forces (Fig. 5a, perturbed condition). In contrast when the cost was low, it corrected less (smaller force, larger hand deviation) as expected under the minimal intervention principle(Todorov & Jordan, 2002). Behaviorally, these error corrections recapitulate the effect of environmental factors shown by Nashed and colleagues(Nashed et al., 2012) where the online correction was modulated by the target width. At the network level, despite the absence of motor output during preparation, some network nodes already encoded task objectives (i.e. varied as a function of the lateral cost parameter). At the population level, preparatory PCs separated as a function of the lateral cost parameter, followed by distinct trajectories during execution (Fig. 5b).

**Figure 5.**
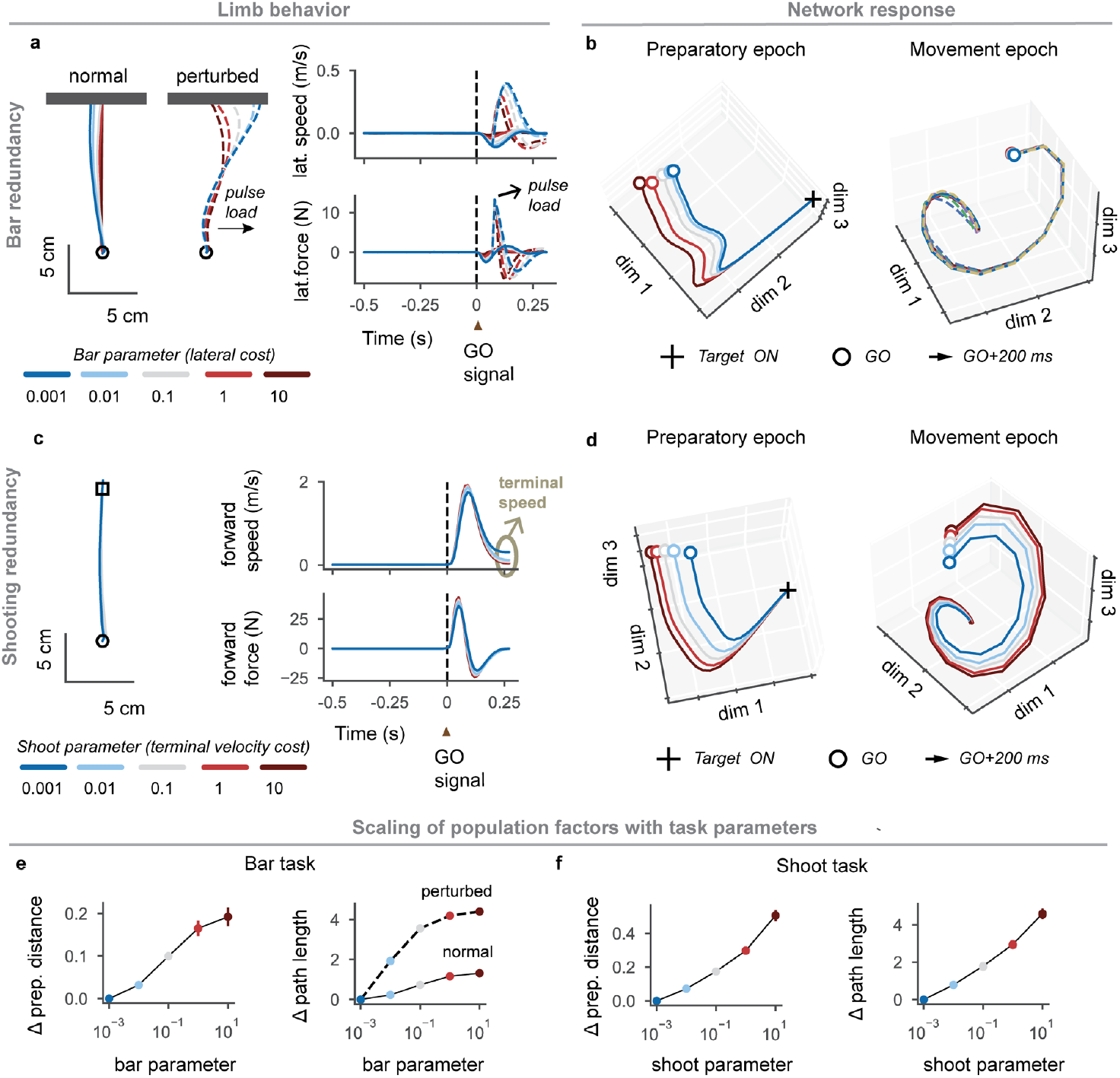
Flexible control to changes in task performance objectives. a) representation of the task with varying levels of penalty on the lateral end-point coordinate allowing modulation of the feedback response projection of the network trajectories on the three principal modes from preparatory and movement epochs for five different levels of lateral terminal displacement cost parameter. (c-d) same as above for five different levels of shooting parameter captured by a cost on termina hand velocity. e) Change in preparatory and movement period factor properties for bar task. f) Same as e for the shooting task. The circle markers depict mean across N=5 networks, and the error bars indicate the s.e.m.

In the shooting task, the penalty on terminal velocity of reaching the target was systematically reduced, leading to shooting movements that hit the target without stopping on target (Fig 5c). Notably, the differences in kinematics were minimal (Fig. 5c), especially the differences in hand speed became apparent only near the end of the movement (Fig. 5c top-right panel). At the population level, preparatory PCs reflected changes in the cost on terminal velocity before the actual movement, followed by distinct movement related PCs (Fig. 5d). Across different network instantiations, both preparatory and movement-related PC trajectories scaled with cost parameters for both tasks, demonstrating that these parameters can be extracted from the low-dimensional features of the network activity (Fig. 5e, f). Movement PCs during perturbation showed larger path length modulation with the bar parameter (Fig. 5e), highlighting how state-feedback control integrates limb feedback to adjust network trajectories according to task objectives.

Interestingly, our model reproduced another unexpected feature of M1 activity linked to task-dependent consideration of limb feedback. We applied constant loads in eight radial directions during a prepare-to-reach task toward a target 20 cm from the start location (Supplementary Fig. 6). Loads were presented unexpectedly in two contexts: 500 *ms* after target onset (preparatory epoch) and 100 *ms* after the GO cue (execution epoch). In both cases, the controller rapidly corrected hand deviations. Simulations revealed that the magnitude of individual nodes’ responses to loading during preparatory (posture) and execution epochs were uncorrelated (Supplementary Fig. 6a, b). Such random changes in the load representations were consistent with evidence of distinct cortical responses during posture maintenance and reach execution(Kurtzer et al., 2005). Note however that our model applied end-point loads in cartesian space, unlike joint torques used in the abovementioned experimental study.

### Low rank hypothesis applied to network-body system

Our developments and simulations highlight a critical role of state-feedback and task parameters in shaping population activity. First, the optimal input to the system clearly conveys feedback from the modelled limb into the network (Eqn. 3), which must influence the network trajectories. The analysis of the network’ activity without feedback also confirmed the defining role of state feedback on the network trajectories (Fig. 3). Next, our previous simulations show that both task parameters and perturbation responses together influence network trajectories (Fig. 5e, Supplementary Fig. 6). Note that we do not specify the source of inputs, in practice biological systems must contend with delays and noise in sensory processing and it is assumed that an estimate from sensory feedback or internal predictions is available(Franklin & Wolpert, 2011; Shadmehr & Krakauer, 2008). Now, we develop a more formal argument of why low-dimensional features emerge from limb feedback based on the spectral properties of the network-body system.

The dynamics of the full system (Eqn. 4) can be written compactly as ż = *A*_*c*_ ***z***, where *A*_*c*_ represents the combined system dynamics of the network, limb and desired limb states under closed-loop control. For simplicity we ignored the constant input (***ε***) as it does not impact subsequent analyses. The matrix *A*_*c*_ has a block structure, consisting of a high-dimensional, partially random component (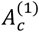) and a low-dimensional component (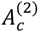) related to the limb feedback gains (Fig. 5a, see Methods).

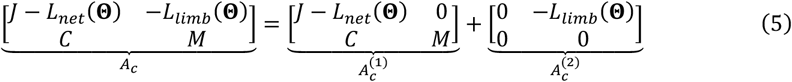

Thibeault and colleagues(Thibeault et al., 2024) recently showed that the connectivity of many natural physical systems can be modelled as a combination of a random component and a low-dimensional component. They used Weyl’s inequality to show such systems are effectively low-dimensional, provided the spectrum of the random component is small in comparison with the low-dimensional component of the overall system connectivity. By rewriting the Weyl’s inequality from(Thibeault et al., 2024) while considering Eqn. 5, we get:

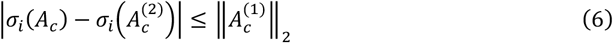

for all *i* ∈ {1,2, …, *n*}, where *σ*_*i*_ is the *i*^*th*^ singular value and ‖. ‖_2_ is the spectral norm (largest singular value) of a matrix. The inequality indicates that, when the spectral norm of 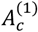 is sufficiently small, the singular values of the total connectivity matrix (*A*_*c*_) remains close to the singular values of its low-dimensional component (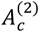)(Thibeault et al., 2024). In this scenario, the total matrix (*A*_*c*_) also attains a low effective rank, meaning that a small number of dominant features capture most of the connectivity structure.

We examined the dimensionality of network structure under the application of feedback control in the center-out reaching task. In line with the low rank property (Eqn. 6), the singular values of the matrix *A*_*c*_ decreased rapidly, and matched those of the limb-feedback connectivity term 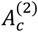 for the first four components (Fig. 6a-b). Using a well-defined effective rank metric that is based on the uniformity of singular value distribution (see Methods), we found that the dimensionality of the network structure closely tracked that of the limb related input. Consequently, even though the system’s initial connectivity was high-dimensional (*erank*: 69), the structure of the controller input forced low-dimensionality (*erank*: 3).

**Figure 6.**
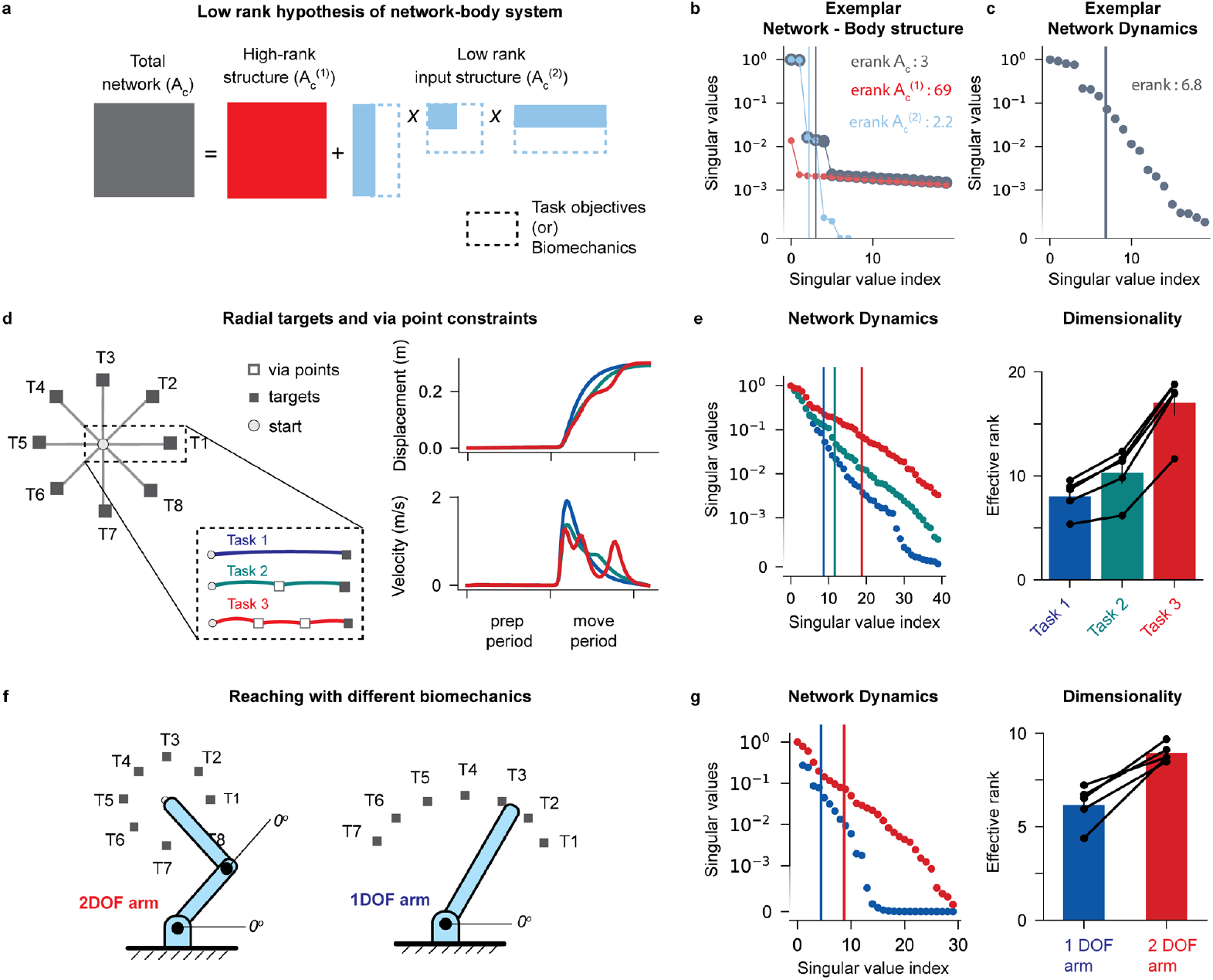
Testing low-rank hypothesis in the interacting network-body system. a) decomposing network structure into the default connectivity and the limb input structure resulting from the structure of the LQR controller. We indicated the low-dimensional structure of 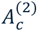 by illustrating the singular value decomposition of this matrix. The dashed lines indicate that the rank of the low-dimensional components can change depending on the task definition and dimension of the effector. See main text (Eqn. 5) for the mathematical description of these block matrices. b) Singular values of the composite matrix (in gray), of the default connectivity (in red), and the limb input matrix (in cyan). c) Singular values of covariance matrix of network trajectories obtained after concatenation of their trajectories across the 8 different targets. Observe that the effective rank (<7) was low in comparison with the network size (n = 100). d) Illustration of the task with via-point used to modulate the complexity (left subpanel). Simulated hand displacement and velocity to a horizontal target in the three via-point tasks (right subpanels). e) Singular values of the network activity in the movement period (left), effective ranks of the covariance matrices of the network trajectories across five different instances of the model. The vertical lines are the effective ranks, the horizontal axis was truncated for clarity. f) Illustration of the eight-directional reaching task with a 2-degree of freedom (2-DOF) and a 1-degree of freedom (1-DOF) arm. g) Singular values of the covariance matrix of the network activities in the movement period (left), and effective ranks for five different instances of the model (right). The light gray lines on the bar plots correspond to individual network instances.

Thus, it is expected that the network activity be also low-dimensional as their dimension is determined by the connectivity structure. Indeed, the effective rank of the covariance of network activity across time and targets was much lower (*erank*: 6.8) than the actual number of network nodes (*n* = 100) (Fig. 6c).

The foregoing spectral analysis connected the rapid decay of the singular values of the system matrix *A*_*c*_ (i.e., its structural properties) to the low-dimensionality of the network activity (i.e., its dynamics). A direct corollary is that increasing the dimensionality of the input signal (through 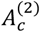) containing task-relevant information should increase the effective rank of the network activity (dashed lines in Fig. 6a). We tested this in two ways: (1) first we increased task complexity by adding via-points along the movement path (2) then we modified the dimension of the plant by considering either a single joint system or a two-joints system.

We simulated the center-out reaching task with 0, 1, or 2 intermediate via-points. Although hand paths were generally similar (Fig. 6d), the velocity profiles showed additional sub-movements as the number of via-points increased. We observed two key changes indicating an increase in the dimensionality of feedback input signals. First, the effective rank of the network connectivity component associated with the limb feedback (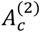) increased with the number of via-points (Supplementary Fig. 7a). Second, the time series of network inputs also showed an increase in effective rank (Supplementary Fig. 7b). Given that the dimensionality of the network activity is tightly linked to the structure of limb feedback, we examined the changes in the singular values of the network activity. As task complexity increased, the singular value spectrum decayed more slowly (Fig. 6e). Consequently, the effective rank, equivalent to the dimensionality of the network activities, increased with task-complexity (Fig. 6e), although the network structure, eight reach directions, and movement duration remained unchanged. This suggest that increasing task-demands selectively induced higher-dimensional feedback input signals, thereby increasing the dimensionality of the associated network dynamics.

We next varied limb biomechanics by simulating movements with either a one-dimensional (1-DOF) or a two-dimensional (2-DOF) limb to reach eight radial targets (Fig. 6f). The number of DOF directly modified the size of the state vector and of the feedback gain matrix *L*_*limb*_ (**Θ**). Again, the effective rank of the network connectivity component associated with limb feedback, and the time series of network inputs increased with the added biomechanical dimension (Supplementary Fig. 7c, d). Singular value decomposition of the network activity showed that the 1-DOF simulations exhibited a steeper singular value decay, resulting in a lower effective rank than the 2-DOF simulations (Fig. 6g). Thus, increasing the biomechanical dimensions also increased the dimensionality of the corresponding network dynamics.

Together, these results support the view that low-rank motor cortical dynamics emerge naturally from optimal control of the network-body system, and provide a formal connection between the dimensionality of the input to the network, such as the number of degrees-of-freedom or the presence of intermediate goals, and the neural population activity. Finally, as a corollary of low rank hypothesis described above, we observed that for similar tasks and biomechanics, and therefore similar inputs to networks, the principal components were preserved across variations in the network connectivity (Supplementary Fig. 8). This observation is consistent with the observed similarity in neural population trajectories across individuals performing a task(Safaie et al., 2023), assuming that different brains can be characterized by a similar network topology.

## Discussion

Our results demonstrate that several features of neural population dynamics naturally emerge from optimal control of a simple, linear network-body system. By coupling a high-dimensional random network to a biomechanical plant and computing optimal control gains for each task, we replicated several hallmark features of motor cortical activity including rotational dynamics(Churchland et al., 2012; Sabatini & Kaufman, 2024), preparatory states encoding the parameters of impending movements(Churchland et al., 2010; Churchland & Shenoy, 2024; Kaufman et al., 2014), and orthogonal preparatory and execution subspaces(Elsayed et al., 2016). Our model captured the fact that population activity reflects variations in task parameters, such as altered force fields(Ahmadi-Pajouh et al., 2012; Sun et al., 2022), inertial load(Kirk et al., 2024), or task demands before movement onset. Finally we demonstrated that this linear formulation establishes the critical role of the input to the network (here the state-feedback control signal) in the shaping of the network dynamics(Sauerbrei et al., 2020) based on the low-rank hypothesis of complex systems(Thibeault et al., 2024).

Clearly, the model was based on very simplifying assumptions: we assumed a fully observable state, direct readout for the actuation signals, and perfect knowledge of the network connectivity to derive the optimal controller. We believe that these assumptions were justified to formulate the basic properties of such a network-body system in the simplest case, and expect that future works unveil the properties of more biologically plausible models in detail. Clearly, considering a partially observable state (i.e., with delays and noise(Scott, 2016)), and indirect connections between the body and the network may enrich the predictions with insight into distributed processes across different brain regions(Michaels et al., 2020, 2025; Omrani et al., 2016) including cerebellum(Becker & Person, 2019; Israely et al., 2025), somatosensory and parietal areas(Chowdhury et al., 2020; Guo et al., 2025; Omrani et al., 2016; Takei et al., 2021), and spinal cord(Guang et al., 2024; Kirk et al., 2024; Lindén et al., 2022). Likewise, the question of robustness against imperfect knowledge of the network structure in the controller is an important challenge for prospective studies.

The model architecture corresponds to assumptions that are consistent with experimental evidence from motor cortical recordings: First, the relationship between network and muscle activation is fixed (Eqn. 1), which is consistent with the evidence that neural activation patterns exhibit a fixed relationship with muscle activation when changing arm posture or adapting to a force field (Cherian et al., 2013; Scott et al., 1997). Likewise, network units exhibited changes in response magnitudes dependent on the task(Kurtzer et al., 2005). However, we did not model the effect of delays and nonlinearities that are known to determine neural responses during reaching and mechanical loading(Lillicrap & Scott, 2013; Pruszynski et al., 2011). These simplifying assumptions are not fundamental limitations since they can be addressed in more complex instances of the network-body model, by modifying the block-structure of the network(Perich et al., 2021), including nonlinearities in the muscle and body dynamics(Lillicrap & Scott, 2013; Trainin et al., 2007), or adding delays and noise following standard techniques(Crevecoeur et al., 2016; Takei et al., 2021). In this light, the simplifying assumptions discussed above can be seen as an asset: it allowed us to comprehensively demonstrate the remarkable explanatory power of the simplest instance of the model, thereby showing that the first basic, linear approximation already provides deep insight along with testable predictions about the shape and dimensions of neural factors that one may expect for any experiment that could be modeled in this framework.

Our approach differs from conventional neural network models which rely on data-driven training(Aldarondo et al., 2024; Chiappa et al., 2024; Codol et al., 2024; Feulner et al., 2022, 2025; Guang et al., 2024; Kalidindi et al., 2021; Lillicrap & Scott, 2013; Merel et al., 2019; Michaels et al., 2016, 2020; Saxena et al., 2022; Sussillo et al., 2015; Zimnik & Churchland, 2021), with task-dependent network trajectories that can be recalled by contextual inputs(Mante et al., 2013) when similar situations arise. Here, because we formulated a linear model, the optimal network weights could be calculated analytically as optimal feedback control gains modifying both the way state feedback was fed into the network, as well as how different network nodes needed to be (de)coupled. This approach offers an original interpretation of how optimal controllers can be synthesized in the brain: variations in model parameters (**Θ**) yield distinct feedback control gains that modify input to the system and coupling of network nodes. Thus, task-selection (e.g. hold then move)(Elsayed et al., 2016; Kurtzer et al., 2005), task-dependent control(Donchin et al., 1998; Gallego et al., 2018; Heming et al., 2019; Michaels et al., 2025; Miri et al., 2017; Mizes et al., 2024; Rodriguez et al., 2024; Saxena et al., 2022), and motor adaptation(Kirk et al., 2024; Perich et al., 2018; Sun et al., 2022) can in principle be achieved in a fixed-network architecture by efficiently modulating the projections of somatosensory signals into the network according to the task parameters.

This view provides a mechanism by which humans can express flexible, task-dependent control across fast timescales, i.e., between or even within trials(Braun et al., 2009; Česonis & Franklin, 2022; Comite et al., 2022; Crevecoeur, Mathew, et al., 2020; Crevecoeur, Thonnard, et al., 2020; Kalidindi & Crevecoeur, 2024; Kobak & Mehring, 2012). While connectivity may be reshaped over longer timescales through synaptic plasticity(Feulner et al., 2025; Gurnani et al., 2024; Ostry & Gribble, 2016; Sadtler et al., 2014), rapid adjustments may be plausibly explained by swiftly modifying feedback control inputs through a fixed network architecture(Schimel et al., 2023; Sohn et al., 2021). We do not rule out that real sensorimotor circuits may develop specialized connectivity for commonly encountered tasks over time(Singh & Scott, 2003), owing to plasticity in cortical and subcortical regions (Pemberton et al., 2024). Rather, we propose that such experience-driven plasticity coexists with fast, task-dependent modulation of feedback-driven input to the network, enabling primates to deploy flexible control policies across a wide range of contexts very rapidly.

Another key insight offered by the linear formulation was that low-dimensional neural factors can emerge because limb and task parameters provide a dominant, low-rank input to the otherwise high-dimensional recurrent network. This aligns with the low-rank hypothesis of complex systems(Thibeault et al., 2024), which posits that a small subset of principal modes embedded within a large random network yields a low effective rank, making the system trajectories low-dimensional. The block structure of the network-body dynamics and the expression of optimal control gains (Eqn. 5) allowed decomposing the closed-loop system into components linked to the network or feedback input, and study the spectral properties of each component as well as their effect on the dimension of the network trajectories. A direct consequence of this relationship was that the dimensionality of neural trajectories scaled with task complexity or with the dimensionality of the effector (Fig. 5, via points and degrees of freedom, respectively). Overall, this model portrays motor cortex as an input-driven dynamical system, consistent with experimental evidence(Israely et al., 2025; Omrani et al., 2016; O’Shea et al., 2022; Park et al., 2025; Sauerbrei et al., 2020; Vahidi et al., 2024), while additionally highlighting the role of state feedback and task parameters in the computation of optimal network inputs.

In all this study shows that linear models as first approximation can provide a powerful test bed to make interpretable predictions about the shape and dimensions of neural trajectories, providing a candidate theoretical link between neural activity and sensorimotor control. This framework predicts that variations in task parameters lead to measurable changes in population activities, reflecting distinct control policies and biomechanical constraints. These normative insights may inform fundamental research and brain-machine development to overcome the challenges posed by increasingly large neural datasets.

## Methods

We used the following convention throughout the paper: scalar quantities appear in italicized symbols, vectors in bold and italicized font, matrices in capital symbols, and sets of parameters in bold and capital symbols. The dependency on time is sometimes omitted for clarity.

### Network dynamics

We generated random networks comprising *n* = 100 units. The network activity was represented by the rate vector ***r***(*t*) denoting the firing rate of all network units. The dynamics of this network was governed by the rate equation:

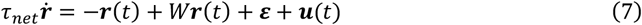

Here *τ*_*net*_ = 20*ms* is the time constant of a single network unit, and *W* is the network adjacency matrix. For compactness, we describe the network dynamics by combining the first two terms into 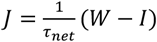, where *I* is the identity matrix. Throughout this article, we refer to *J* as the network state transition matrix. The input to the network consists of two terms: ***ε*** ∈ ℝ^*n*^ is a constant vector drawn from a zero-mean normal distribution 𝒩(0, 5) generating an offset representing spontaneous activity. This constant offset was treated with system augmentation considering 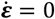. The vector ***u***(*t*) is the control signal that must be calculated based on the cost function representing task demands. In our model, we assumed that all units receive direct control inputs. We observed that restricting direct control inputs to a subset of network units produced similar results. The network adjacency matrix *W* was initialized such that all eigenvalues of the resulting state transition matrix *J* were negative. We studied three classes of random networks described below.

The first class of networks consisted of *Inhibition Stabilized Networks (ISN’s)*, comprising separate inhibitory and excitatory populations, with *n*_*E*_ = 80 excitatory units and *n*_*I*_ = 20 inhibitory units. The combined network activity was represented by the rate vector 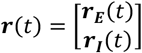, where ***r***_***E***_(*t*) and ***r***_***I***_ (*t*) denote the firing rates of the excitatory and inhibitory populations, respectively. To simulate inhibitory stabilized regime, the network adjacency matrix was constructed using the algorithm described by Hennequin and colleagues in (Hennequin et al., 2014; Kao et al., 2021).

The second class of networks consisted of *Fully Connected Recurrent Networks*. Here, the network adjacency matrix was drawn from a normal distribution 𝒩(0, *g*^2^/n), where g = 0.8. This configuration ensured stable recurrent dynamics, as described previously(Michaels et al., 2020; Sussillo et al., 2015).

The third class of networks corresponded to *Erdös-Rényi* random networks, in which connections between unit pairs were determined probabilistically. Specifically, each connection was assigned with a given probability parameter, with *p* = 0.4 indicating that only 40% of total number of neuron pairs were connected (other parameter values yielded comparable results). Excitatory and inhibitory weights were assigned by randomly setting nonzero connections to +0.5 or −0.5 with equal probability. To ensure network stability, we first set the diagonal terms in the adjacency matrix to zeros. This was followed by incremental decrease in the diagonal terms until all eigen values became negative.

Overall, the given network drives limb movement by actuating muscles through an output vector ***a***(*t*). The relationship between network activity and limb muscle actuation was defined as:

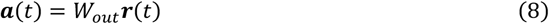

Here *W*_out_ ∈ ℝ^2×*n*^ is the readout matrix whose entries are drawn from a normal distribution, 𝒩(0, 0.5), allowing multiple network units to contribute at different amounts to limb actuation. Notably, each random network corresponded to different random samples of the adjacency matrix, while maintaining a fixed readout matrix.

### Limb dynamics

For the point mass approximation of limb dynamics, the limb state vector was defined as 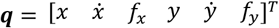 where the coordinates *x* and *y* represent the displacement of point mass in the lateral and forward directions respectively, the dot represent their time derivatives, and *f*_*x*_ and *f*_*y*_ are the forces applied by the controlled actuator to the mass. These forces respond to the network’s output ***a***(*t*), with *a*_*x*_ and *a*_*y*_ representing the network’s actuation in the *x* and *y* directions, respectively. Overall, the system of differential equations for the limb and the actuation dynamics was:

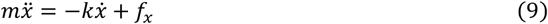

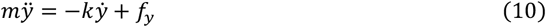

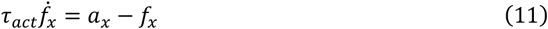

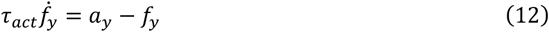

Here *k* is the dissipative constant (0.1 Ns*m*^−1^), and Eqn (11), (12) represent the first order muscle dynamics with the actuation time-constant *τ*_*act*_ = 0.1 *s*.

The limb state evolves according to the differential equation 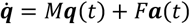, with:

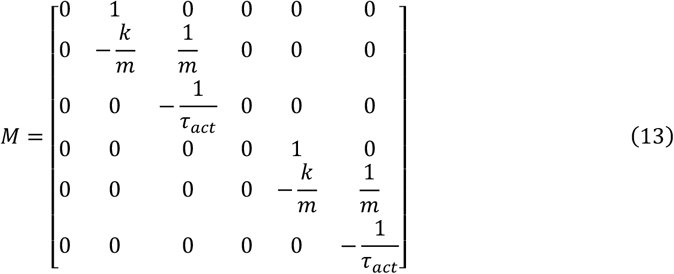

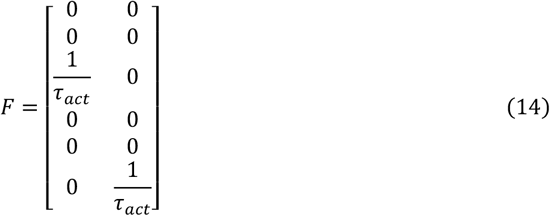

Additionally, we have simulated a linear approximation of a two-dimensional planar arm actuated at the shoulder and elbow joints. Details on the methods and simulations with this shoulder-elbow arm are provided in the Supplementary Material.

For simplicity, we can define a joint variable 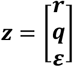 to jointly describe the limb and network states, including the constant offset term for spontaneous activity, expressed as 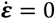. The dynamics of this joint system can be expressed compactly as ż = *A****z*** + *Bu*, where:

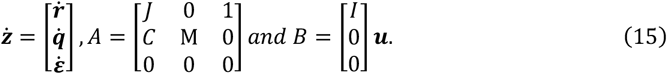

Here 0 and 1 represents a block matrix filled with 0’s and 1’*s* respectively, and *I* represents an identity matrix with size equal to the number of network units. Here, *C* = *FW*_out_ represents the transformation (*FW*_out_) from network activity (***r***) to the forces acting on the limb (***f***). Note the matrix *A* here indicates the default state-transition matrix of the network-body system. Applying closed-loop control, as follows, involves modifications in this state-transition matrix with feedback components.

### Receding horizon control in a Linear Quadratic Regulator (LQR) form

We solved the problem of finding optimal network control inputs by repeatedly minimizing a cost on limb states and inputs within a moving time horizon

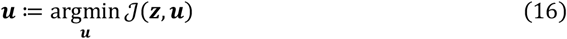

Given the quadratic form of the cost function (Eqn. 2), and the linear system dynamics (Eqn. 3), the optimal control reflects the well-known LQR problem (Åström, 1970; Todorov, 2005), whose solutions takes the following state-feedback form

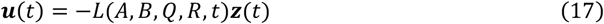

Here, *L* is a time-varying control gain matrix that linearly maps the available state measurements (***z***) or their estimates onto control inputs.

To compute the control gains, we transformed the control problem into a discrete time system based on Euler integration with 10 *ms* time steps. Then the control gains were computed recursively backward in time following standard techniques (Åström, 1970; Todorov, 2005). This procedure yields control gains that depend on the system dynamics (*A* and *B*) and on the task objectives (*Q* and *R*), as represented in Eqn. 17. Calling **Θ**^(*i*)^ = *A*^(*i*)^, *B*^(*i*)^, *Q*^(*i*)^, *R*^(*i*)^ the set of task parameters known to the controller for a given task (*i*), we can rewrite the control law as:

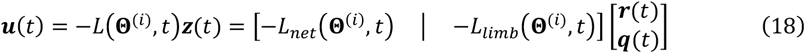

This formulation expresses that the state-feedback control policy is specific to a given task parameter set **Θ**. Notice that in results section, we denoted the task parameter set with individual block components (*J, C, M*) of the system matrix (*A*), but they are essentially the same. Notably, the control law can be decomposed to separate the contribution of network or limb related feedback.

Overall, the Receding Horizon Control scheme (Guigon, 2023) iteratively recalculates the optimal control gains at each step. This process involves:

1. Collecting the task parameters **Θ**^(*i*)^ corresponding to the current task (*i*) at each time step (*t*)
2. Calculating the control gains over the prediction horizon {*L*(**Θ**^(*i*)^, *t*), …, *L*(**Θ**^(*i*)^, *t* + Δ*h*)}, for the current time window (from *t* to *t* + Δ*h*).
3. Applying the first control gain of the sequence, ***u***(*t*) = *L*(**Θ**^(*i*)^, *t*)***z***(*t*)
4. Updating the time to (*t* + 1) and repeating from step 1.

In this work, we assumed that the knowledge of task parameters is available and expect that adding a module to perform inference about these parameters when they are not known exactly may link our developments to more general contexts (Heald et al., 2021).

### Prepare-to-reach task

This task involved holding posture at a given starting location (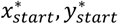) for an unspecified duration, then executing the movement towards the target (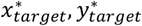) upon receiving a “GO” signal. This paradigm effectively induces a transition between postural control (prepare) and reaching (move) control following a signal which occurs at a non-predictable time. To capture the fact that the occurrence of the “GO” signal was unpredictable, we employed a two-phases receding horizon cost-function with three terms as follows:

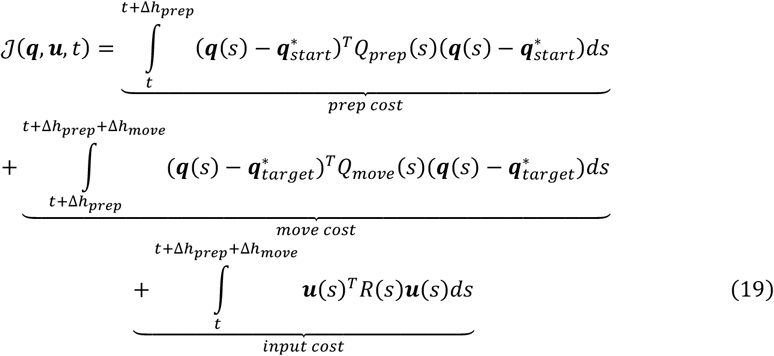

Here, 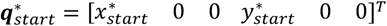 and 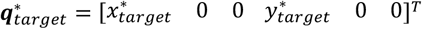 represent the overall desired start and terminal target hand states. Note that the velocity and force terms of the desired states are set to 0, meaning any deviations from the non-zero velocities and forces incurs a cost. Here, *Q*_*prep*_(.) and *Q*_*move*_(.) penalize state deviations during the preparatory and movement epochs, respectively. Before the “GO” signal, the first term penalizes deviations from the starting position over a predicted preparatory horizon Δ*h*_*prep*_ = 50 *ms*. This horizon maybe much shorter than the actual preparatory period (typically > 500 *ms*), to encourage rapid preparation for the uncertain GO signal. The second term penalizes deviations from the target over a fixed movement horizon Δ*h*_*move*_ = 500 *ms*, starting at *t* + Δ*h*_*prep*_. The third term penalizes control inputs over both horizons. This cost was recomputed at each time step during the preparatory period until the “GO” signal occurs.

Once the GO signal was delivered, the predicted preparatory horizon was replaced by the measured time of the GO signal (*t*_*GO*_). The cost function for the movement period became:

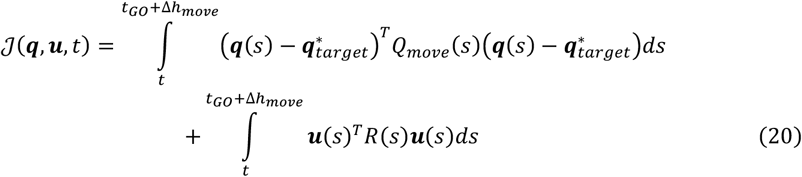

The first term penalizes the deviations from the target state, while the second term penalizes control inputs over the fixed movement horizon. By expressing the distinct objectives of preparatory and movement periods into separate cost terms, this optimization naturally yields two distinct sets of feedback control gains: one for maintaining posture while anticipating a movement during preparation, and another for executing the reach once the GO signal arrives.

We use the following parameter values, *Q*_*prep*_ = *diag*([100,100 0, 0,100,100,0]) for the cost on hand deviation from the desired start location, and *Q*_*move*_ = *diag* ([10, 0.1, 0,10,0.1,0]) for the cost on hand deviation from the desired terminal target location. The terminal target cost penalty *Q*_*move*_ was further scaled by the factor 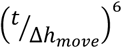 to ensure low initial cost when the elapsed time (*t*) is small relative to the movement horizon (Δ*h*_*move*_), and a smooth build up until the terminal time step. The penalty on the network control inputs was set at *R* = 10^−10^ ensuring the solutions prioritize reaching the targets with relatively smaller cost on the squared magnitude of control inputs.

### Changes in Limb Mechanics

We investigated two specific scenarios: a velocity-dependent force field and an inertial loading task. In the Velocity-Dependent Force Field Task (FF), the limb was subjected to a velocity-dependent force field where the force *f*_*x*_ was proportional to the effector’s velocity 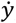, expressed as 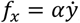, and the force *f*_*y*_ was proportional to the effector’s velocity 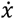, expressed as 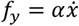 as in standard motor adaptation paradigms. This introduced a coupling between forward velocity – lateral force, and lateral velocity – forward force, effectively modifying the limb dynamics matrix *M* to *M*(*α*). The limb dynamics including the force fields can be represented as follows:

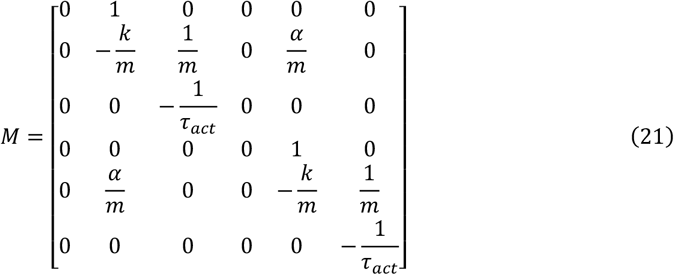

In the Inertial Loading Task, we simulated the addition of inertial loads (*m*_*load*_), which altered the limb dynamics. These additional loads influence the limb dynamics matrix *M*, resulting in a modified matrix *M*(*m*_*load*_). This allowed us to examine how the control scheme adapted to changes in limb inertia. The limb dynamics with the additional inertial loads was formulated as follows:

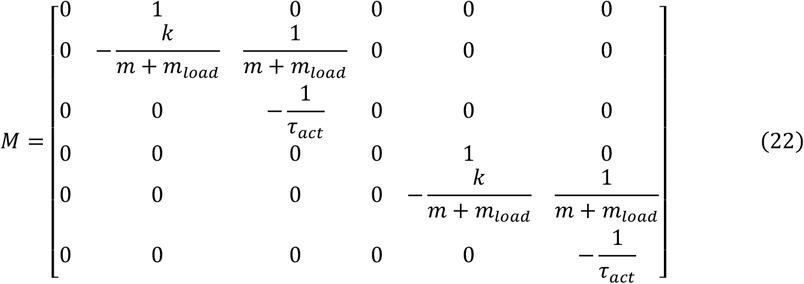

We simulated the prepare-to-reach task using the receding horizon control scheme across five different levels for each parameter: *α* ∈ {−5, −2.5,0,2.5,5} *NSm*^−1^ and *m*_*load*_ ∈ {0.2, 0.4, 0.6, 0.8, 1.0} *KG*. Notably, the cost parameters were the same across the three simulated reaching tasks: center-out reaching, force field and inertial loading.

### Changes in Task Objectives

To investigate the influence of task objectives on the network-body dynamics, we modulated matrix *Q* that assigns cost penalty on each state component.

In the spatial target redundancy task, we focused on the lateral deviation to simulate reaching to a target with different geometry (as the dot-bar task (Nashed et al., 2012)). We systematically varied the parameter governing lateral deviation in the cost matrix *Q*_*move*_ from Eqn. 19-20 by multiplying the default value with a scalar *λ*_*bar*_ across five levels: {0.001, 0.01, 0.1,1,10}. This scalar multiplication essentially transforms the matrix penalizing state deviation as *Q*_*move*_ = *diag*([100 × *λ*_*bar*_, 0.1, 0,100,0.1,0]). A low parameter value (e.g., when multiplied with 0.001) imposed minimal penalty on lateral (*x*) deviation, allowing for larger deviation in endpoint location. Conversely, higher parameter value (e.g., when multiplied with 10) imposed a strong penalty on the lateral deviation, necessitating corrective actions to achieve a low terminal displacement.

In the “shooting” task, we focused on the hand velocity. We varied the cost associated with the lateral and forward velocity parameters by multiplying the default value with a scalar *λ*_*shoot*_ across the same five levels: {0.001, 0.01, 0.1,1,10}. This scalar multiplication transforms the terms penalizing the hand velocity by transforming the matrix *Q* _*move*_ = *diag* ([100, 0.1 × *λ shoot*, 0,100,0.1 × *λ* _*shoot*_, 0]). At low parameter value (e.g., when multiplied with 0.001), stopping at the target was not required, resulting in non-zero terminal velocity—consistent with a “shooting” motion. At higher values (e.g., when multiplied with 10), terminal velocity was heavily penalized, necessitating deceleration or stopping of the hand as it reached the target.

In both bar and shooting tasks, the penalty on the network control inputs remained unchanged at *R* = 10^−10^.

Both biomechanical adaptation (modifying matrix *M*) and flexibility in task objectives (adjusting matrix *Q*) involved changes in the task parameter set **Θ**^(*i*)^. As described in Eqn. (18), alterations in these parameters modified the control gains *L*(*t*), leading to task-dependent network-body trajectories. By systematically varying the parameters associated with limb dynamics (*M*) and the entries in the cost function (*Q*), we assessed how the control system adapted its policy to maintain optimal performance across different biomechanical conditions and task objectives.

### PCA in preparatory and movement epochs

Following previous work by Elsayed and colleagues (Elsayed et al., 2016), we grouped the network responses into two matrices corresponding to the preparatory and movement epochs. Preparatory epoch responses were represented by a rectangular matrix *P* ∈ ℝ^*n*×*c*d^, where *n* is the number of units, *c* is the number of conditions (e.g., number of targets, force-field values, bar target penalties, via-point scenarios etc.,), and *d* is the total duration of the preparatory epoch. Similarly, movement execution period responses are represented by *E* ∈ ℝ^*n*×*cd*^. We first soft-normalized the activity by the activity range plus a small constant (0.005). We performed PCA on the covariance matrix of preparatory response epochs *P*, treating each row (i.e., network unit) as a variable, to obtain the top six components (prep-PCs) of the network trajectories in the *n*-dimensional space. Similarly, we obtained the top six components from the movement epochs (move-PCs) by performing PCA on the covariance matrix of movement response *E*.

We compared how the pairwise correlations changed between the preparatory and movement epochs by computing the correlation in firing rates between each possible pair of units during each epoch. We then computed an average of these correlations across eight target conditions. This yielded two correlations for each pair of units, one for each of the preparation and movement epoch (Fig. 2c). We tested if the correlation coefficient for each pair of neurons in one epoch linearly correlated with that of the other epoch. Pearson correlation coefficient was used to quantify the pairwise correlations between the preparatory and movement epochs across all neuron pairs (Fig. 2d).

### Alignment Index

We determined how aligned the top-ten preparatory and movement related principal components were by the alignment index, following the procedure described previously in (Elsayed et al., 2016) :

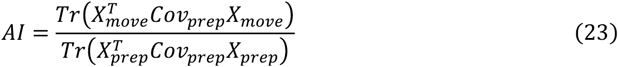

Here *X*_*prep*_ and *X*_*move*_ ∈ ℝ^6×*cd*^ are matrices containing the top six preparatory and movement principal components, *Cov*_*prep*_ is the preparatory covariance matrix, and *Tr*(.) is the trace operator. Eqn. 23 described the ratio of how much variance the top six movement principal components could explain of the preparatory activity with how much top six preparatory principal components could explain of the preparatory activity (the maximum preparatory variance captured by any six linear projections). The alignment index ranges from 0, indicating the preparatory and movement activities are orthogonal, to 1, indicating the preparatory and movement activities are perfectly aligned.

In Fig. 2, we compared the alignment indices of the model with two publicly available monkey M1 recordings from prepare to reach tasks (Churchland et al., 2012; Suresh et al., 2020). The public dataset from (Churchland et al., 2012) predominantly captures execution-related neural activity, with only a brief 50 ms window preceding the onset of the phasic M1 response, that may reflect the states achieved at the end of preparatory epoch. Accordingly, we defined proxy of the preparatory response (*X*_*prep*_) from the 0 − 50 *ms* interval prior to the phasic response onset, and the execution response (*X*_*move*_) from the 50 − 100 *ms* interval following movement onset, ensuring symmetric 50 *ms* windows for both responses. For the model and M1 recordings from (Suresh et al., 2020), which include more extensive preparatory epochs, we defined the preparatory response in the 200 *ms* window spanning 50 − 250 ms before the GO cue. The execution response was defined in the 200 *ms* window from 50 − 250 *ms* following movement onset.

### jPCA analysis

We performed jPCA analysis on the network activity using procedure developed in (Churchland et al., 2012). First, we obtained the top six movement period principal components *X* (6 × *cd* dimensions). We numerically calculated the derivative of *X* yielding 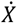, and fit a linear dynamical model which found a relationship between *X* and 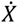, expressed as 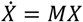. Here, *M* was constrained to be skew-symmetric, which restricted the possible dynamical systems to systems with oscillatory or rotational dynamics. We assessed the model’s fit by calculating the coefficient of determination or goodness-of-fit metric (*R*^2^). jPCA analysis was also applied on publicly available M1 recordings from (Churchland et al., 2012; Suresh et al., 2020). Notably, from (Suresh et al., 2020), we used the data available from Monkey 4 that performed prepare to reach task. Across datasets, jPCA analysis was applied for the first 300 *ms* after movement onset.

### Quantifying rotations and decay in data

To quantify the rotational structure within the neural population activity, we implemented the gyration number analysis proposed by (Kuzmina et al., 2024). This model-agnostic method measures the rotational and decay components present in time-series data without requiring dimensionality reduction. The analysis was performed using the publicly available code from the original publication. The core steps were as follows:

First, neural activity data of movement execution epochs from all experimental conditions were concatenated into a single matrix *E* ∈ ℝ^*n*×*cd*^, where rows represent the number of nodes and the columns represent the activity across *c* = 8 reaching directions and *d* time duration (from 0 to 300 ms after the GO cue). To make the data formatting suitable with the libraries developed by Kuzmina, we transposed the original matrix to *Y* = *E*^*T*^ so that the rows of *Y* represent the time points across all reach directions (*cd*) and columns represent individual nodes (*n*).

Next, we computed spatial differential covariance matrix as 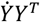, and identified its *i* eigen values (*λ*_*i*_). The coordinates for the gyration plane were derived from the eigen spectrum. The decay axis, which captures expansive or contractive dynamics, and the rotation axis, which captures rotational strength, were calculated using the first complex-conjugated eigenvalue pair as follows:

Rotation axis:

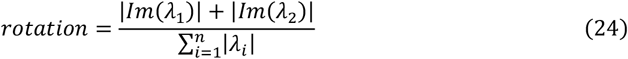

Decay axis:

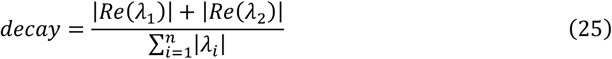

This provided a standardized, two-dimensional measure to compare the rotational dynamics across datasets.

### Tensor Maximum Entropy (TME)

We tested our findings – specifically, low alignment indices, quantity of rotations, and preserved PCs across networks – against the hypothesis that these features are a byproduct of tuning and smoothness properties of neurons. We employed the TME method to generate surrogate datasets (Elsayed & Cunningham, 2017). TME method generates surrogate datasets that retain upto the second-order marginal moments of the original network data (mean, covariances across neurons, conditions, and time) while randomizing the higher-order interaction terms. We generated surrogate datasets for each network across simulated tasks.

In the center-out reach simulated task, we computed the PCs of 1000 randomly sampled TME surrogate datasets, for each network activity dataset, during both the preparation and movement epochs, then calculated the alignment index and the fit to rotational dynamics. We compared these surrogate alignment indices and rotations with those obtained from the actual network data (Fig. 2, Supplementary Fig. 2, 3, 4). Similarly, surrogate datasets were computed for M1 recordings (Fig. 2), and the alignment indices and rotations were compared with those obtained from the actual data.

Furthermore, for each task, we computed the PCs of 1000 randomly sampled TME datasets per network and examined the correlations among these PCs across networks performing the same task (Supplementary Fig. 8).

### Singular value decomposition analysis

We performed singular value decomposition to analyze the dimensionality of the network connectivity matrices and population activities. Recall that the network-body dynamics can be jointly expressed as a closed-loop system of the form presented in Eqn. 4. For simplicity, we can ignore the spontaneous activity offset ***ε***, and the overall system can be represented as follows,

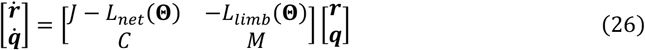

The above network-body dynamics jointly express closed-loop system of the form ż = *A*_*c*_ ***z***, where *A*_*c*_ has a block structure corresponding to network and limb dynamics as well as their connection. Note that *A*_*c*_ represents the closed-loop state-transition matrix that includes control gains, and it is different from the open-loop state-transition matrix *A* described in Eqn. 15 before applying the feedback control law. This matrix was then decomposed into two terms (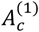and 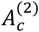) as described in Eqn. 5.

Singular value spectra have been found for each of the decomposed terms and of the composite matrix (Fig. 6a, b). Additionally, singular value spectrum has also been computed for the network responses matrix in the movement epoch, *E* ∈ ℝ^*n*×*cd*^ (Fig. 6c). Here *c* = 8 describes the number of target directions in the center-out reaching task for both the via-point (Fig. 6d) and biomechanical tasks (Fig. 6f).

### Dimensionality

Given the singular value spectra, *σ*_*i*=1,2,…,*n*_, of a matrix *A*, its dimensionality can be measured from the well-defined Roy and Vetterli’s effective rank as described in (Thibeault et al., 2024):

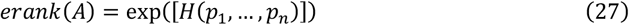

Here *H*(*p*_1_, …, *p*_*n*_) = − ∑_*i*_ *p*_*i*_ *logp*_*i*_ is the Shannon entropy of the singular value spectrum, measured as a function of the singular value mass function

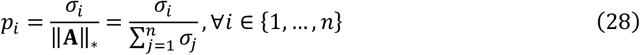

*erank* measures the uniformity of the singular value distribution, resulting in a low effective rank when the singular values decrease rapidly.

## Acknowledgements

HTK was supported by the Fonds de la Recherche Scientifique (FRS-FNRS) Chargé de recherche Grant CR 252 (FC 043127). FC was supported by the FRS-FNRS Grant 1.C.033.18 (FC 036239).

## Author Contributions

Conceptualization: H.K and F.C; Data curation: H.K; Formal analysis: H.K; Funding acquisition: H.K and F.C; Investigation: H.K; Methodology: H.K and F.C; Project administration: H.K and F.C; Resources: F.C; Supervision: F.C; Validation: H.K and F.C; Visualization: H.K; Writing – original draft: H.K and F.C; Writing – review & editing: H.K and F.C.

## Code Availability

The code and data from simulations will be available in the public repository ‘figshare’ after publication

