## Supplementary Material for "A model of neural population dynamics for flexible sensorimotor control"

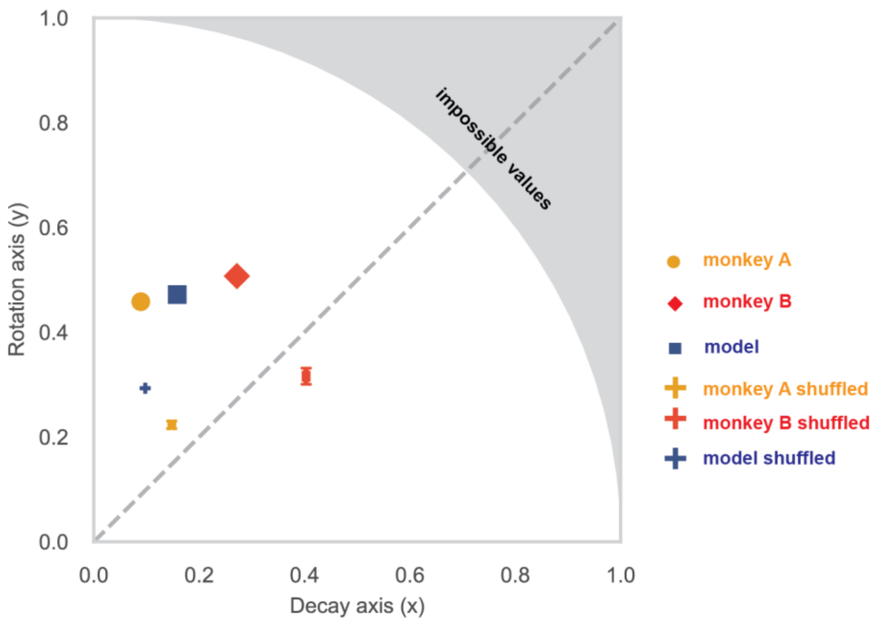


***Supplementary Figure 1 Analyzing the rotations and decay in the rate responses using procedure developed by Kuzmina and colleagues (Kuzmina et al., 2024).*** *Monkey A data is from Churchland et al., 2012, and Monkey B is from the dataset of Monkey 4 in Suresh et al., 2020. The shuffled datasets were generated using the open code repository made available by Kuzmina and colleagues that replicated the procedure used in Churchland et al. 2012. The error bars represent* $\pm SEM$

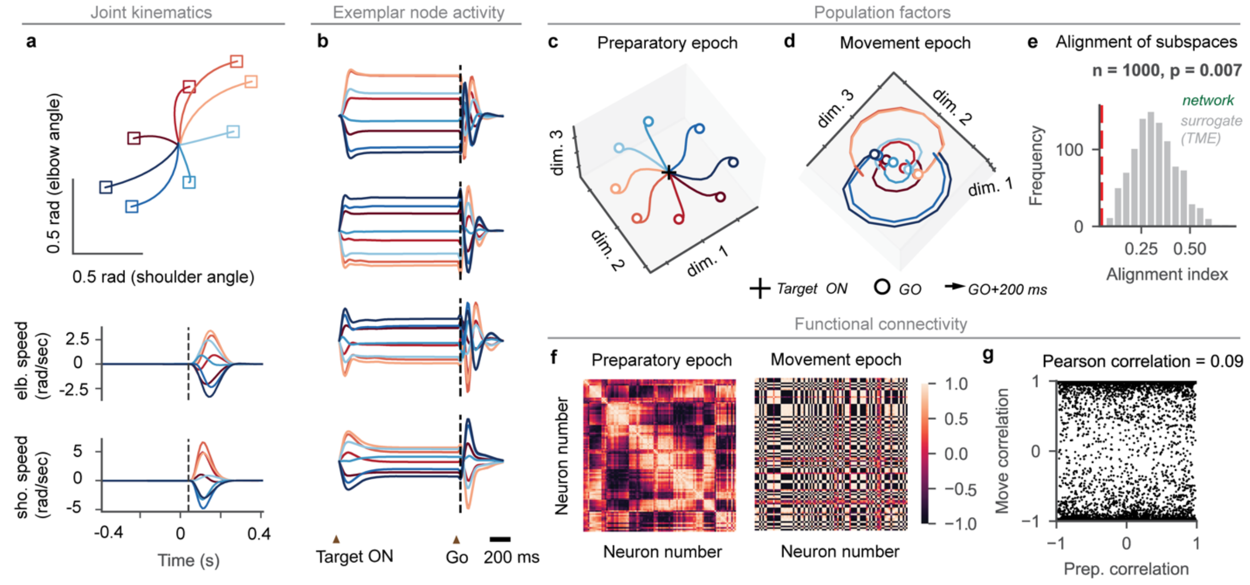


***Supplementary Figure 2 Simulations of center-out reaching on a Linearized planar 2-DOF arm.*** *a) Joint kinematics for the simulation of a shoulder-elbow arm. The arm dynamics was linearized around the starting joint configuration* $(30°, 90°)$*, and the state space was described in terms of joint states* $\boldsymbol{\theta} = [\theta_1, \theta_2, \dot{\theta}_1, \dot{\theta}_2, \zeta_1, \zeta_2]^T$ *representing joint displacement, velocity and torques at the shoulder and elbow joints respectively (see 2-link planar arm dynamics section in the Supporting Information). b) exemplar neural firing rates from selected units, observe that the activity of some neurons depends on the target of the pending movement c) neural activity along the principal modes in the preparatory period d) same as c for the movement period e) alignment between preparatory and movement period subspaces. The histogram shows distribution of alignment indices across surrogate data and the vertical line represent the alignment of subspaces from the principal modes of preparatory and execution epochs (low alignment corresponds to orthogonal). f) Pairwise correlations in preparatory and movement periods g) Linear relationship between pairwise-correlations from movement to preparatory period. The lack of pattern shows that activities from one epoch is statistically independent of that of the other epoch.*

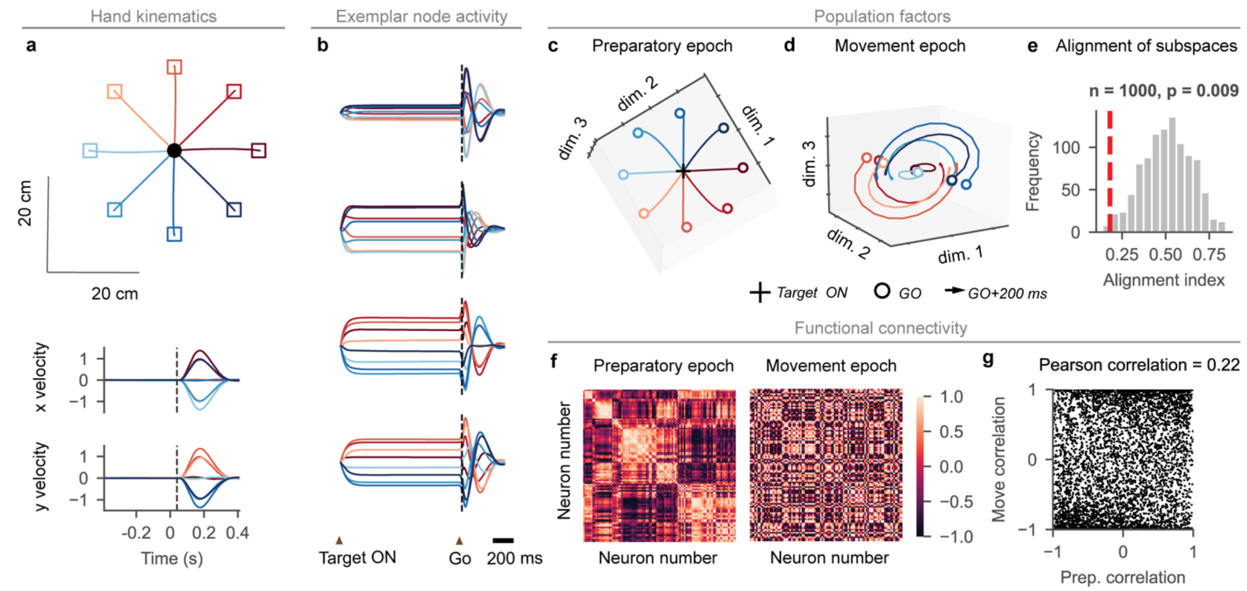


***Supplementary Figure 3 Simulations of center-out reaching using a fully connected RNN with random connectivity.*** *a) Hand kinematics for the simulation of a point-mass system b) exemplar neural firing rates from selected units, observe that the activity of some neurons depends on the target of the pending movement c) neural activity along the principal modes in the preparatory period d) same as c for the movement period e) alignment between preparatory and movement period subspaces. The histogram shows distribution of alignment indices across surrogate data and the vertical line represent the alignment of subspaces from the principal modes of preparatory and execution epochs (low alignment corresponds to orthogonal). f) Pairwise correlations in preparatory and movement periods g) Linear relationship between pairwise-correlations from movement to preparatory period. The lack of pattern shows that activities from one epoch is statistically independent of that of the other epoch.*

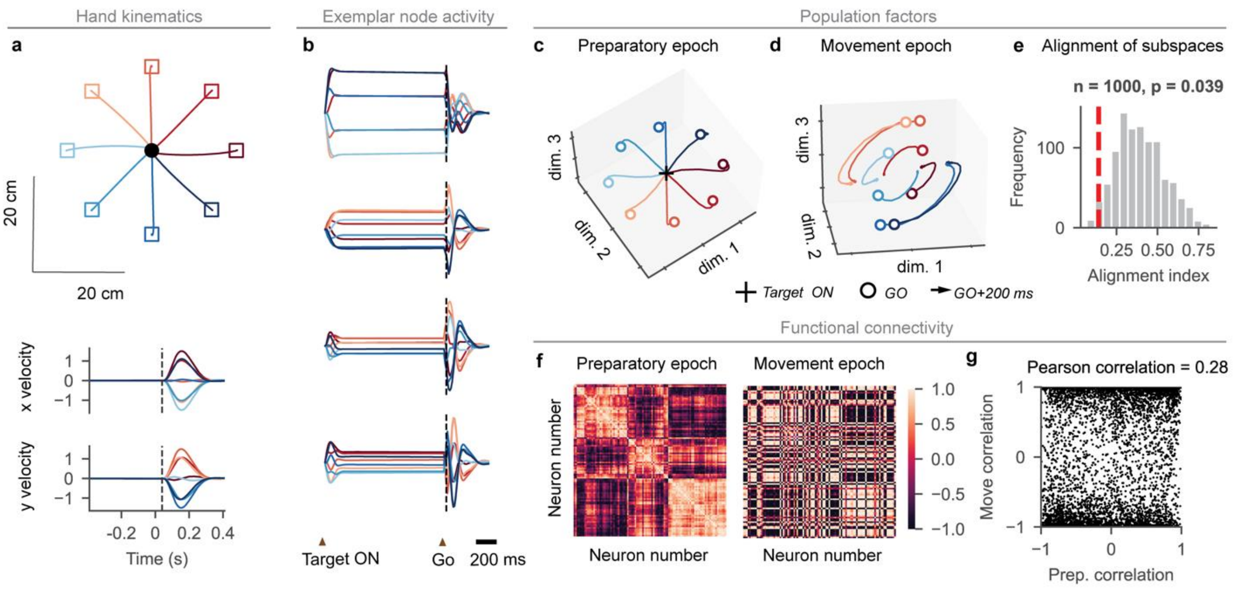


***Supplementary Figure 4 Simulations of center-out reaching using an Erdös-Rényi connected network.*** *a) Hand kinematics for the simulation of a point-mass system b) exemplar neural firing rates from selected units, observe that the activity of some neurons depends on the target of the pending movement c) neural activity along the principal modes in the preparatory period d) same as c for the movement period e) alignment between preparatory and movement period subspaces. The histogram shows distribution of alignment indices across surrogate data and the vertical line represent the alignment of subspaces from the principal modes of preparatory and execution epochs (low alignment corresponds to orthogonal). f) Pairwise correlations in preparatory and movement periods g) Linear relationship between pairwise-correlations from movement to preparatory period.*

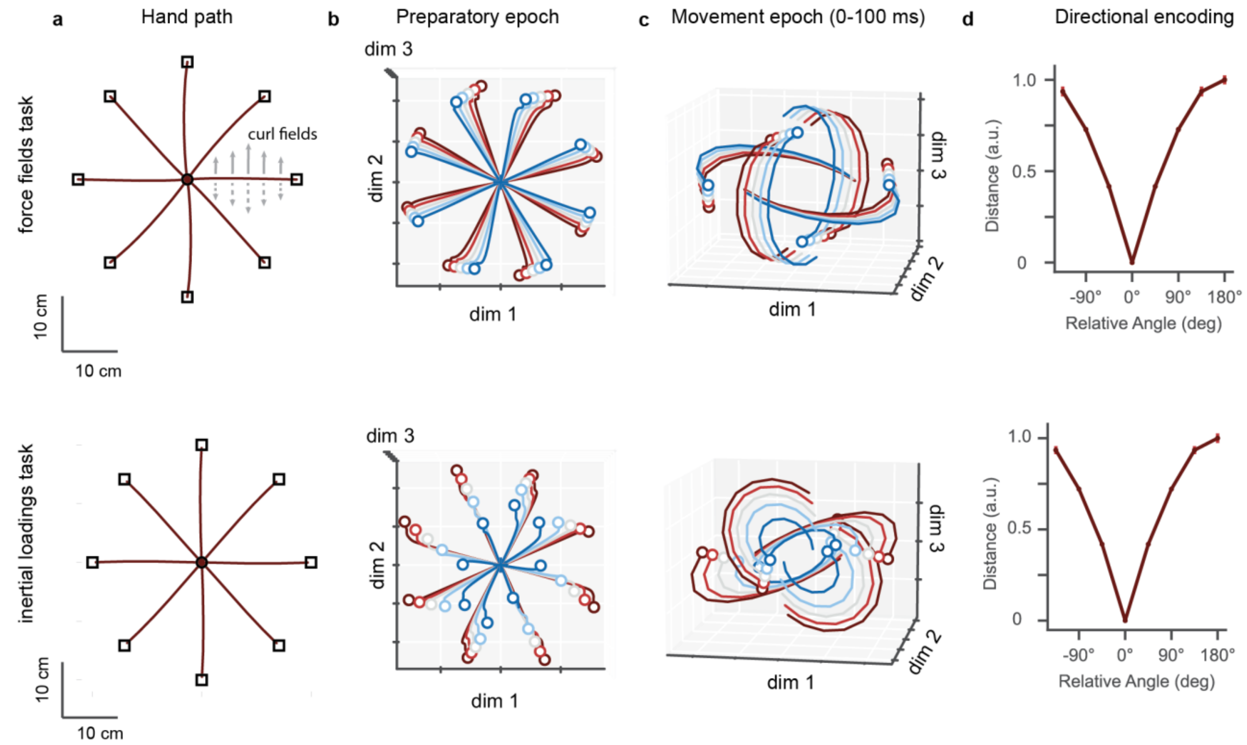


***Supplementary Figure 5 Simulations of center-out reaching with mechanical loads*** *a) Hand path towards 8 targets, across 5 different levels of velocity-dependent force fields or inertial loads applied moving towards each target b) first three preparatory period PCs from eight directional targets and different levels of variations in movement loading c) same as b for execution PCs. Note that we plotted only 4 out of 8 target directions in the execution PCs to simplify the visualization without a clutter d) Distances computed as root mean square differences between preparatory states of a specific reach direction from all the other reach directions and normalized by the maximum distance across reach directions. We computed this value by making each direction of reach a reference point and compared against all the other directions. The plot shows mean and standard deviation across n=5 different networks.*

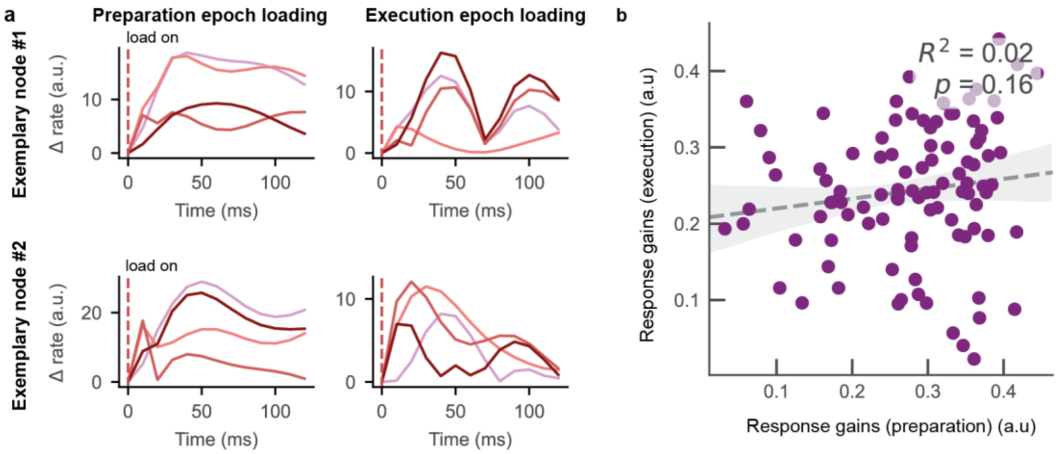


***Supplementary Figure 6. Load sensitivity of network nodes during preparation (posture) and execution (reaching) epochs.*** *a) change in activity of exemplary nodes from a no load condition, after sudden mechanical loading using a constant load. Each color trace corresponds to a different direction of load. The activity represents the absolute value whereas the original traces can be both positive and negative. Each color corresponds to a different direction of load (between 0 – 180 degrees) b) Posture epoch gains were computed by regressing node activity of n=100 nodes in the preparatory period in 0-100 ms time window after perturbation onset against the load directions. Reach epoch gains were computed by regressing node activity between 0-100 ms after perturbation onset in the movement period against the load directions. Each point corresponds to the response gains of a single network node in posture and movement phases. The analyses procedure was similar to previous experimental study Kurtzer et al., 2005.*

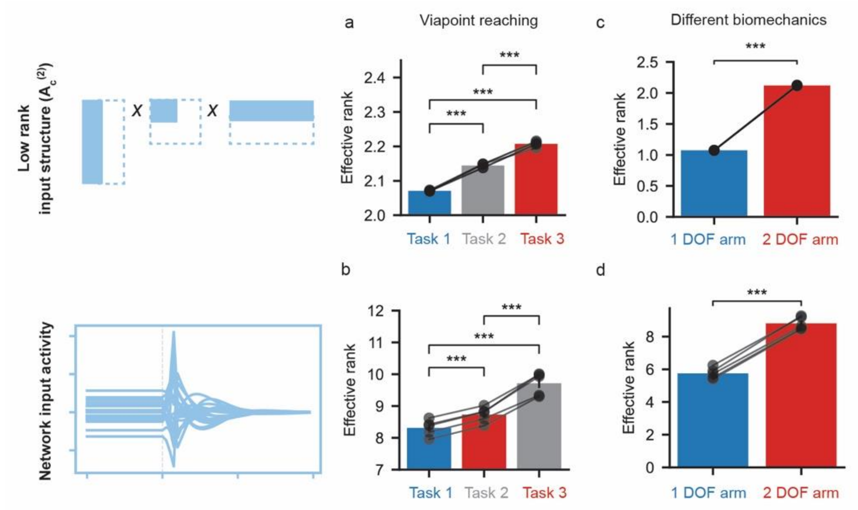


***Supplementary Figure 7 Dimensionality analysis of the input feedback signals.*** *a) effective rank of the low rank input structure across via point tasks with 1,2 and 3 targets b) effective rank of the covariance matrix of the network input signals in the via point reaching tasks c) effective rank of the low rank input structure across the limb variations d) effective rank of the covariance matrix of the network input signals across limb variations.*

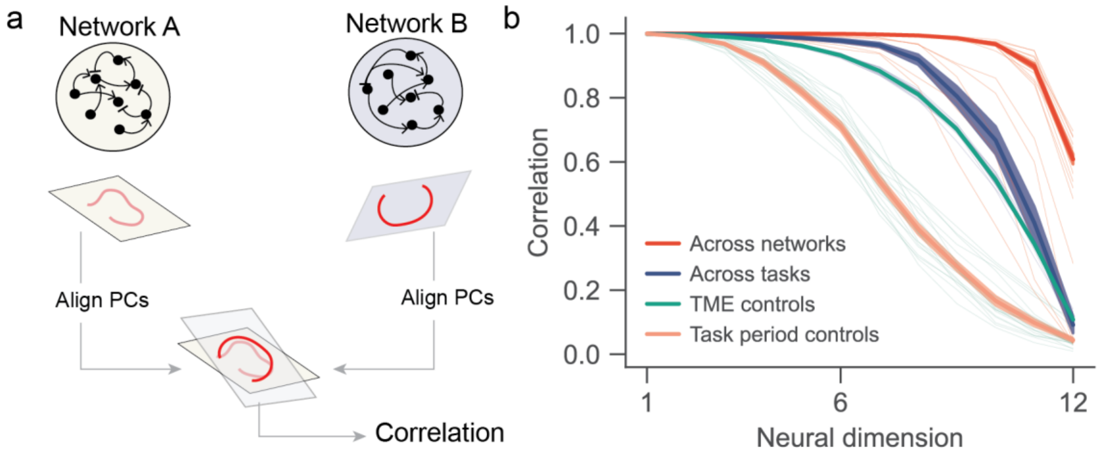


***Supplementary Figure 8 Preserved latent dynamics across networks performing the same task on the same limb.*** *a) schematic illustration of the two-step procedure used to compare PCs across networks. First, we used canonical correlation analysis (CCA) to find the appropriate linear transformations of the top 12 PCs of each network for maximum alignment. Next, we computed the Pearson correlations of each of the 12 aligned PCs and ranked them in a descending order. b) Ranked correlations of the aligned PCs. 'Across networks' indicates correlations found across networks (n=5) performing the same task (for each of the 4 tasks: force field, inertial load, bar and shoot tasks) on the same biomechanics. When task and biomechanics were fixed, the aligned PCs were highly correlated across networks. 'TME controls' indicate the correlations between matched surrogate data generated from the original network data in all tasks. 'Task window controls' indicate the correlations found with original network data, but where the task periods were randomized (i.e., PCs of two networks were from different time periods within the task duration). Both control datasets yielded significantly lower correlations than the actual data. 'Across tasks' indicates the correlations in PCs between all different pairs of tasks for a given network, measured for each of the (n=5) networks. Networks performing different tasks exhibited lower correlations compared to the task-fixed condition, suggesting the consistency of task related inputs influence how preserved the principal components are to network variations. Thick lines are the mean values, and the shaded regions indicate the standard errors. Thin lines are individual correlations computed at each pair of datasets (networks, tasks, and controls)*

**2-link planar arm dynamics**

Here we describe the forward dynamics of a two-joint limb, with torque actuated shoulder and elbow joints. that Let $\boldsymbol{\theta} = \begin{bmatrix} \theta_1 \\ \theta_2 \end{bmatrix}$ represent the joint angles of shoulder and elbow respectively. The forward dynamics of the two-joint limb has been described as follows:

$$\ddot{\boldsymbol{\theta}} = \mathcal{M}(\boldsymbol{\theta})^{-1}\big(\boldsymbol{\zeta} - \mathcal{B}\dot{\boldsymbol{\theta}}\big) \tag{1}$$

$$\mathcal{M}(\boldsymbol{\theta}) = \begin{pmatrix} a_1 + 2a_2\cos(\theta_2) & a_3 + a_2\cos(\theta_2) \\ a_3 + a_2\cos(\theta_2) & a_3 \end{pmatrix} \tag{2}$$

$$\mathcal{B} = \begin{pmatrix} b_{11} & b_{12} \\ b_{21} & b_{22} \end{pmatrix} \tag{3}$$

where $a_1 = I_1 + I_2 + m_2{l_1}^2$, $a_2 = m_2 l_1 c_1$ and $a_3 = I_2$; $l_1 = 0.145\ m$ and $l_2 = 0.284\ m$ are the lengths; $m_1 = 0.2108\ kg$ and $m_2 = 0.1938\ kg$ are the masses; $I_1 = 0.025\ kg.m^2$ and $I_2 = 0.045\ kg.m^2$ are the inertias; $c_1 = 0.0749\ m$ and $c_2 = 0.0757\ m$ are the distance between the end and the center of mass of the two segments. Furthermore, $b_{11} = b_{11} = 0.1$ and $b_{21} = b_{12} = 0.025$. For simplicity, we eliminated the Coriolis term that exerts velocity dependent coupling of shoulder and elbow joints.

The joint torques $\boldsymbol{\zeta} = [\zeta_1, \zeta_2]^{\boldsymbol{T}}$ follow a simple first order dynamics

$$\tau_{act}\dot{\boldsymbol{\zeta}} = -\boldsymbol{\zeta} + \boldsymbol{u}$$

where $u$ is the external input, and $\tau_{act}$ is the time constant for torque actuation. We linearized the arm dynamics around origin of the total limb state $\big[\theta_1, \theta_2, \dot{\theta}_1, \dot{\theta}_2, \zeta_1, \zeta_2\big]^T = [30°, 90°, 0,0,0,0]^T$, resulting in the following limb dynamics:

$$\begin{bmatrix} \dot{\theta}_1 \\ \ddot{\theta}_1 \\ \dot{\zeta}_1 \\ \dot{\theta}_2 \\ \ddot{\theta}_2 \\ \dot{\zeta}_2 \end{bmatrix} = \begin{bmatrix} 0 & 1 & 0 & 0 & 0 & 0 \\ 0 & \dfrac{b_{21} - b_{11}}{a_1 - a_3} & \dfrac{1}{a_1 - a_3} & 0 & \dfrac{b_{22} - b_{12}}{a_1 - a_3} & \dfrac{-1}{a_1 - a_3} \\ 0 & 0 & -\dfrac{1}{\tau_{act}} & 0 & 0 & 0 \\ 0 & 0 & 0 & 0 & 1 & 0 \\ 0 & \dfrac{a_3 b_{11} - a_1 b_{21}}{a_3(a_1 - a_3)} & \dfrac{-1}{a_1 - a_3} & 0 & \dfrac{a_3 b_{12} - a_1 b_{22}}{a_3(a_1 - a_3)} & \dfrac{a_1}{a_3(a_1 - a_3)} \\ 0 & 0 & 0 & 0 & 0 & -\dfrac{1}{\tau_{act}} \end{bmatrix} \begin{bmatrix} \theta_1 \\ \dot{\theta}_1 \\ \zeta_1 \\ \theta_2 \\ \dot{\theta}_2 \\ \zeta_2 \end{bmatrix} + \begin{bmatrix} 0 & 0 \\ 0 & 0 \\ \dfrac{1}{\tau_{act}} & 0 \\ 0 & 0 \\ 0 & 0 \\ 0 & \dfrac{1}{\tau_{act}} \end{bmatrix} \boldsymbol{u}$$

Finally, the mapping from joint coordinates ($\boldsymbol{\theta}$) and velocities to the hand position and velocity in cartesian space ($\boldsymbol{q}$) is accomplished via forward kinematic mapping

$$\boldsymbol{q}(t) = \begin{bmatrix} x \\ y \end{bmatrix} = \begin{bmatrix} l_1\cos(\theta_1) + l_2\cos(\theta_1 + \theta_2) \\ l_1\sin(\theta_1) + l_2\sin(\theta_1 + \theta_2) \end{bmatrix}$$
